# Spatial integration during active tactile sensation drives elementary shape perception

**DOI:** 10.1101/2020.03.16.994145

**Authors:** Jennifer Brown, Ian Antón Oldenburg, Gregory I. Telian, Sandon Griffin, Mieke Voges, Vedant Jain, Hillel Adesnik

**Affiliations:** Department of Molecular and Cell Biology, University of California, Berkeley; The Helen Wills Neuroscience Institute

## Abstract

Active haptic sensation is critical for object identification and manipulation, such as for tool use in humans, or prey capture in rodents. The neural circuit basis for recognizing objects through active touch alone is poorly understood. To address this gap, we combined optogenetics, two photon imaging, and high-speed behavioral tracking in mice solving a novel surface orientation discrimination task with their whiskers. We found that orientation discrimination required animals to summate input from multiple whiskers specifically along the whisker arc. Many animals discriminated the orientation of the stimulus *per se*, as their performance was invariant to the specific location of the presented stimulus. Two photon imaging showed that populations of neurons in the barrel cortex encoded each of the discriminated orientations, and this coding depended on integration over the whisker array. Finally, acute optogenetic inactivation of the barrel cortex strongly impaired surface orientation discrimination, and even cell-type specific optogenetic suppression of layer 4 excitatory neurons degraded performance, implying a role for superficial layers in this computation. These data suggest a model in which spatial summation over an active haptic array generates representations of an object’s surface orientations. These computations may facilitate the encoding of complex three-dimensional objects during active exploration.

## Introduction

In order to judge the shape, size and location of objects, cortical circuits must integrate information arriving from different parts of the sensor array, whether this be from different parts of the retina, the body surface, or the cochlea. Many animals, including humans, move their sensors to optimize information gathering, a process termed ‘active sensation’ (Gibson, 1962). Humans use active haptic sensation to investigate and manipulate objects by executing various stereotyped hand motions (Lederman and Klatzky, 1987). Tactile input from adjacent fingers convey multi-dimensional information about the orientation and structural relationship of the object’s surfaces to ultimately give rise to a coherent object percept. In the primate somatosensory cortex, individual neurons can encode the orientation and curvature of a stimulus (Hsiao et al., 2002)(DiCarlo and Johnson, 2000). Sensing the shape of an object, however, depends on dynamic integration of tactile input (i.e., skin indentation) with proprioceptive information about hand and joint position (Hsiao, 2008). How cortical circuits extract shape information during active sensation is poorly understood. It remains unclear whether and how neural circuits integrate input from the active haptic array to encode shape.

The rodent vibrissal system is a powerful model for active sensation and tactile perception, and the mouse offers the genetic tools that make dissecting the cellular and circuit basis of tactile perception possible (Petersen, 2019). Much as primates use their hands, rodents sweep their whiskers across objects to localize and identify them (Diamond et al., 2008)(Bush et al., 2016). Judging object location and shape requires sensory circuits to integrate ex-afferent signals due to whisker contact and re-afferent signals due to self-generated whisker motion, conceptually analogous to proprioceptive input from the hand. Subsets of neurons in the barrel cortex integrate these two afferent streams to accurately localize stimuli in a head-centered coordinate frame (Curtis and Kleinfeld, 2009). Presumably similar kinds of computations in the barrel cortex are critical for essential behaviors such as navigation and prey capture. Rodents can readily identify and discriminate objects with their whiskers purely based on shape (Anjum et al., 2006)(Polley et al., 2005), and barrel cortex neurons are sensitive to the location (O’Connor et al., 2010)(Sofroniew et al., 2014)(Curtis and Kleinfeld, 2009), texture (Arabzadeh et al., 2003)(Isett et al., 2018)(Chen et al., 2013), shape (Anjum et al., 2006), angle (Bruno et al., 2003)(Andermann and Moore, 2006)(Lavzin et al., 2012), and motion direction (Vilarchao et al., 2018)(Jacob et al., 2008)(Kwon et al., 2018) of tactile stimuli. Discriminating some, but not all of these tactile features, requires spatial summation over the whisker array (Pluta et al., 2017)(Brumberg et al., 1996)(Krupa et al., 2004)(Kelly et al., 1999). However, despite this work, the neural basis for shape perception during active sensation is not understood. In trained rodents, neurons downstream of the barrel cortex in the posterior parietal cortex can encode the orientation of a stimulus in either the visual or somatosensory modality (Nikbakht et al., 2018). These and prior behavioral results imply that the computation and encoding of surface orientation during active touch may occur upstream in the primary somatosensory cortex or earlier.

Surface orientation is an elementary feature that must be encoded by somatosensory neurons to ultimately give rise to shape perception. To investigate its neural circuit basis, we developed an active surface orientation discrimination task for head-fixed mice, explored the behavioral and sensory basis of task performance, and measured neural activity in the barrel cortex as the mice solved the task. Mice quickly learned to discriminate small changes in orientation using their whiskers. Surface orientation discrimination required integration over vertically stacked whiskers, as performance dropped to near chance levels when the whiskers were trimmed to a single row, but was largely unaffected when trimmed to an arc. Many mice discriminated the surface orientation *per se*, since their performance was unaffected when presenting the oriented stimulus at various horizontal locations around the head. Population calcium imaging in trained mice revealed that neurons in the barrel cortex encoded all presented orientations of the stimulus, a feature that was strongly reduced when mice were trimmed to one whisker. Inactivating the barrel cortex optogenetically strongly reduced performance, suggesting an involvement of cortical circuits. Even much more selective optogenetic inactivation of layer 4 (L4) excitatory neurons reduced task performance. These data help define the circuits and computations that are required for the discrimination of surface orientations during active sensation. The detection and synthesis of multiple surface orientations of an object into a coherent percept of shape are likely to be fundamental to how rodents and humans use touch to identify and manipulate objects for essential behaviors including tool use, prey capture, and food consumption.

## Results

### Mice can use active touch to discriminate surface orientations with high acuity

To probe the neural circuit basis of shape perception, we developed a novel task that required mice to discriminate the surface orientation of a presented stimulus bar (Figure 1A-B). A bar was presented unilaterally to the mouse’s right whisker field in one of eight possible orientations. Mice were trained to discriminate positive angles (‘GO’) and negative angles (‘NOGO’), licking to a waterspout for rewarded ‘GO’ stimuli, but withholding licking to unrewarded ‘NOGO’ stimuli, analogous to previous paradigms that probed active tactile detection (Figure 1C, Supplemental Figure 1) (O’Connor et al., 2010). Trials were self-initiated by the mouse locomoting over a set run threshold, and cued by the audible motion of the stepper motor holding the stimulus bar. Mice routinely reached high levels of performance (Figure 1E-G, I), performed high numbers of trials (301 ± 89 std, n=25 mice, 262 sessions) and reached threshold performance after roughly a week of training (8.3 ± 2.2 std days, n=25 mice, Figure 1H). Mice readily discriminated angles as small as 7 degrees (Figure 1E-G), although it is possible that further training could reveal even finer acuity. We developed a novel photo-interrupt touch detector to precisely quantify the timing and the number of contacts the mouse made before its perceptual decision (Figure 1C, Supplemental Figure 2). The mean latency to lick from the first touch was 0.43 ± 0.07 seconds (std, Figure 1D, n=3 mice, 6 sessions Supplemental Figure 2E) and varied with the stimulus; shorter latencies for more extreme orientations and vice versa, as expected (Supplemental Figure 2E). The average number of touches prior to licking was 15.8 ± 3.6 (std., Supplemental Figure 2) implying that well-trained mice readily solved the task within just a few whisk cycles. We observed a tight coordination between whisking, running and licking behavior in trained but not naïve mice: all trained mice decelerated upon presentation of the stimulus, most strongly for GO stimuli, and simultaneously adjusted their whisker set point caudally along the whisking axis, in a manner strongly dependent on the stimulus orientation (Supplemental Figure 3). Naïve mice showed almost no change in run speed, only a slight reduction in whisker set point, and no licking behavior. These data indicate that multiple motor systems (running, whisking, and licking) were highly coordinated as a consequence of learning in the task.

**Figure 1.**
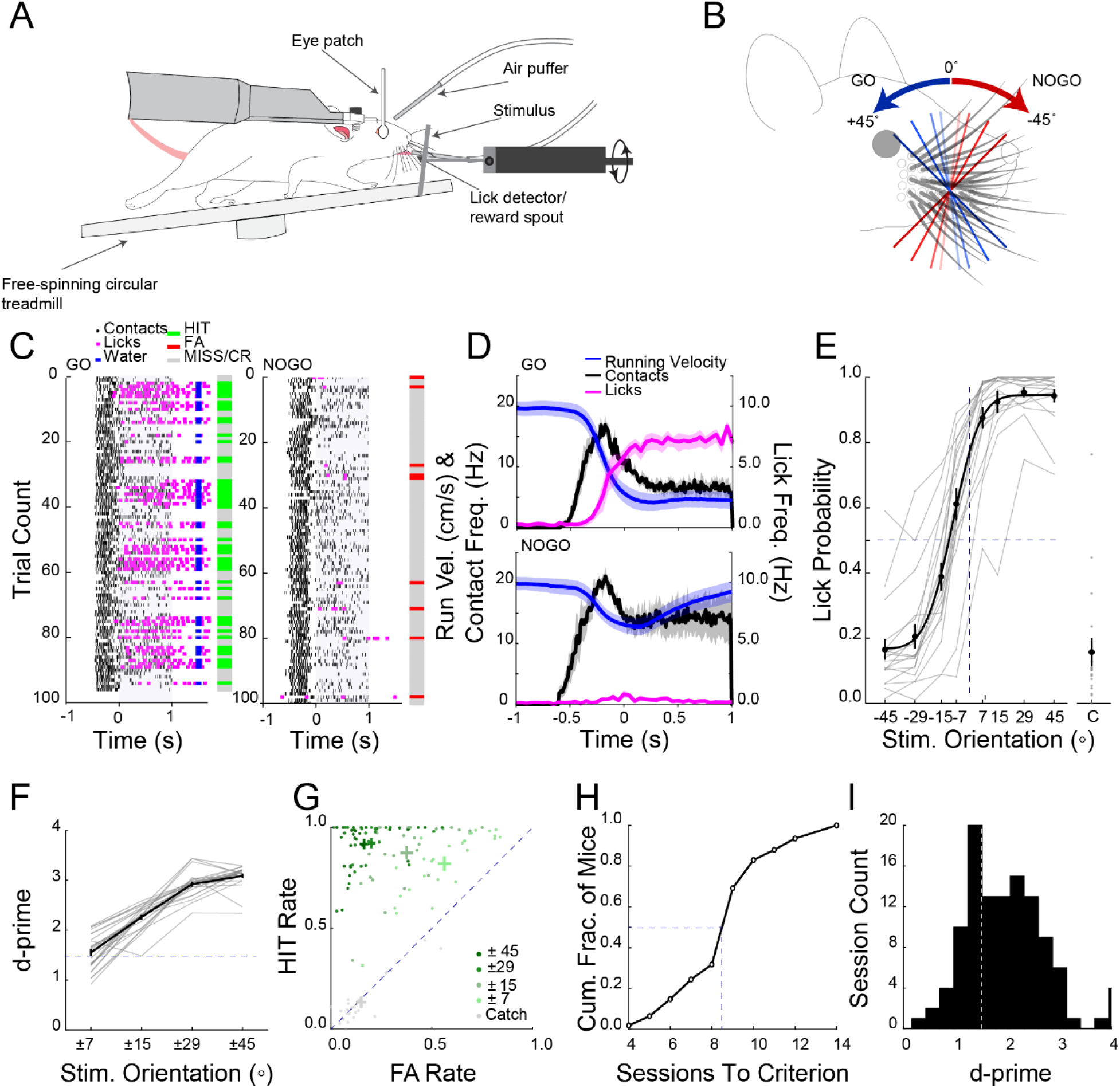
An orientation discrimination task for mice during active haptic sensation. A. Schematic of the orientation discrimination task. Trials are initiated by the mouse running over a run threshold. An orientated bar is presented unilaterally to the mouse’s whisker field. B. ‘GO’ angles (blue shades) and ‘NOGO’ angles (red shades) are pseudo-randomly selected for each trial. C. Raster plot of behavior during a single example session with mouse performing >1.5 d-prime separated into ‘GO’ trials (left) and ‘NOGO’ trials (right). The whiskers contact the stimulus bar (black dots) as the bar moves into the whisking field. Time point 0 represents when the bar stops within the whisker field. Magenta ticks indicate mouse licks, blue ticks show when a water reward was given. Bars to right of raster plots show performance of mouse on each trail (green; ‘Hit’, red; ‘FA’, grey; ‘CR’ or ‘Miss’). D. Average running velocity (blue), whisker contact frequency (black) and lick frequency (magenta) separated into ‘GO’ trials (top) and ‘NOGO’ trials (bottom) (n=3 mice, 6 sessions). E. Psychometric curve of performance (n=25 mice, averages from individual mice (grey lines); group mean ± sem (solid dots and a psychometric fit to group data, solid colored line)). Dashed lines represent midpoints for 0^⍰^ stimulus orientation and 50% lick probability. F. d-prime for corresponding pairs of orientations. G. HIT and FA rates for all sessions over criterion (d-prime>1.5) separated into angle pairs (±45, ±29, ±15, ±7 and ‘catch’ trials). H. Cumulative histogram of the number of sessions to reach criteria (n=25 mice). I. Distribution of d-prime values during task (d-prime 2.3 ±1.2 std, n= 26 mice, 262 sessions). Dotted line represents 1.5 d-prime threshold.

**Figure 2.**
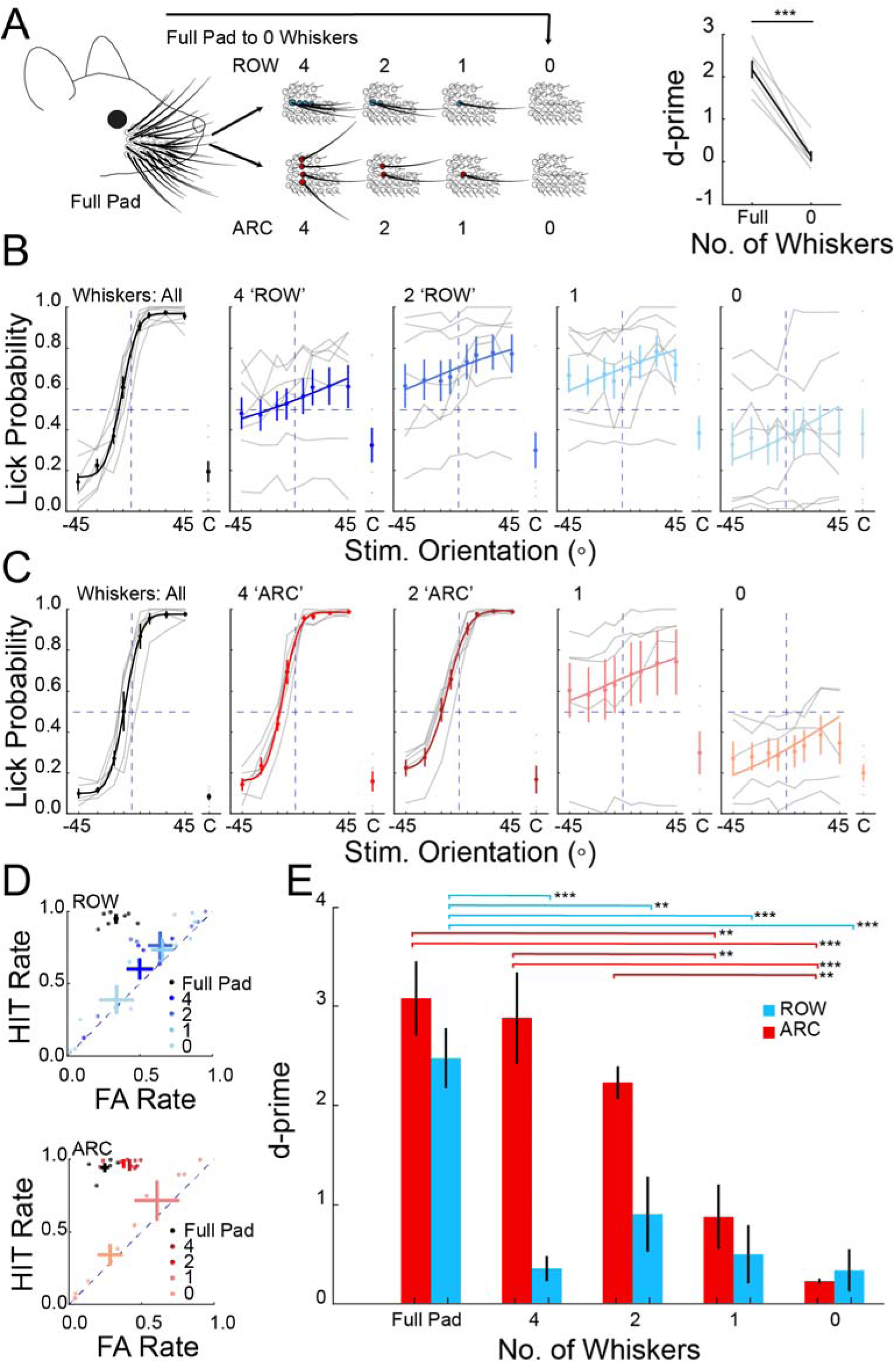
Mice must integrate sensory input from at least two vertically stacked whiskers to discriminate orientations. A. Left: Schematic of the progressive whisker trimming experiment. All mice were trained and reached performance criterion on the task with the full whisker pad. Two separate cohorts of mice underwent progressive whisker trimming to either an arc or a row of whiskers. Right: performance of a third cohort that was abruptly trimmed from full pad to 0 whiskers (p<0.001, n=6 mice, rank sum test). B. Task performance in the cohort of mice in which whiskers were trimmed to a row (n=7 mice; averages from individual mice (grey lines); group mean ± sem (solid dots and a psychometric fit to group data, solid colored line)). C. As in B) but for the cohort of mice trimmed to an arc of whiskers (n=5 mice). D. HIT and FA rate for mice trimmed to a row of whiskers (top) or an arc (bottom). Color shade indicates the number of remaining whiskers in each condition. E. Summary plot of average performance of each group during the progressive trimming paradigm (*p<0.05, **p<0.01, ***p<0.001, Anova).

**Figure 3.**
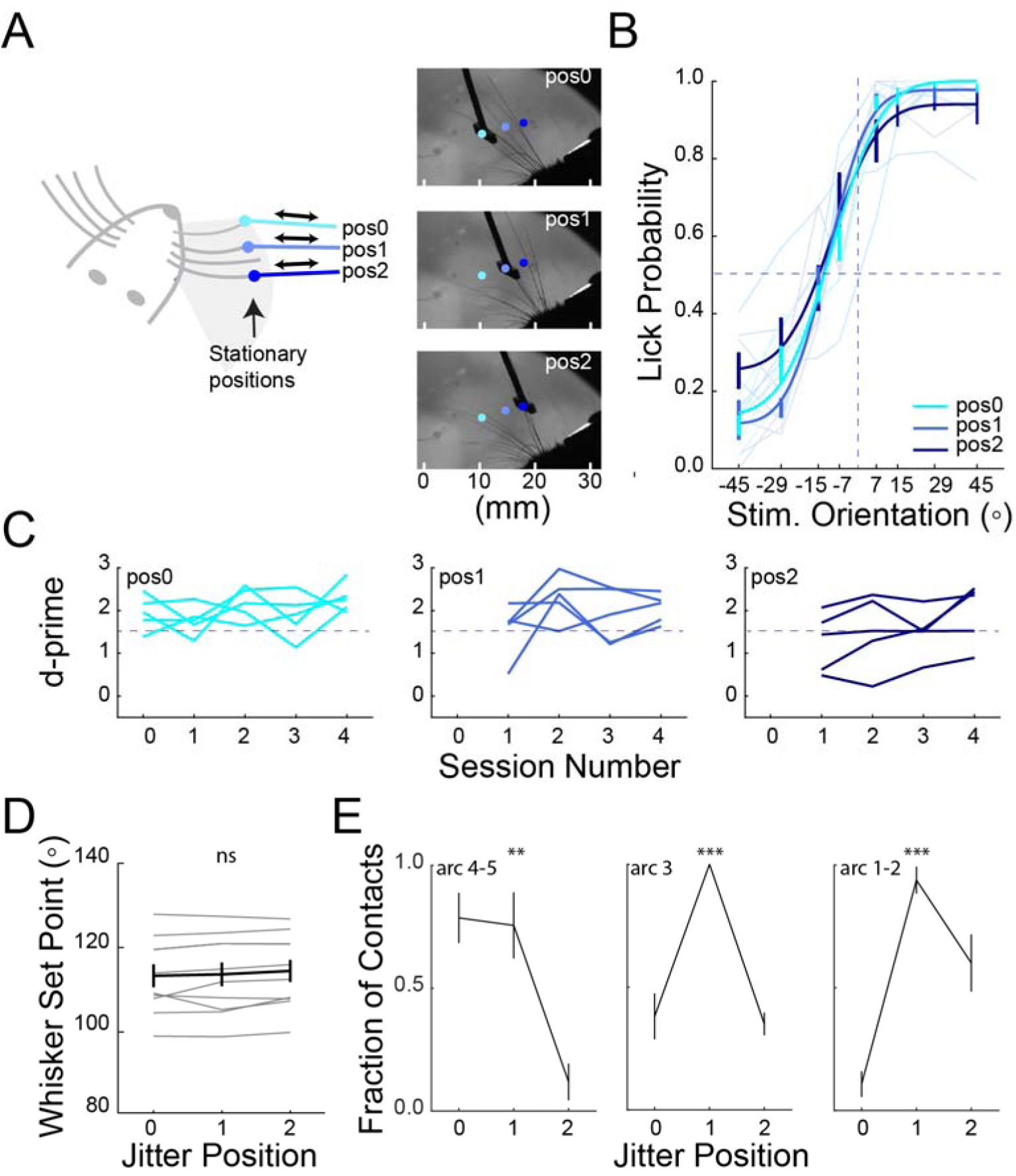
Mice can generalize the orientation discrimination rules to multiple stimulus positions. A. Left: schematic of positional jitter experiment. The stimulus bar was presented laterally to one of 3 different final stationary positions. Right: example images of whiskers contacting stimulus bar in three different jitter positions, from most rostral (pos0, top), middle (pos1, middle), to most caudal (pos2, bottom). B. Average psychometric curves from day 3-4 of jitter experiment for all mice (n=5 mice). Different shades of blue indicate performance data from each of the three final stimulus positions. C. Individual d-prime values for each mouse one day before transition (day 0) and 4 days after introducing linear jitter (day 1-4) separated into each jitter position. Dotted line represents d-prime of 1.5. D. Mean whisker set point across each of the jitter positions (p=ns, Anova). E. Behavioral video analyses to identify the sector (left; arc 4-5, middle (arc 3), right; arc 1-2) of whiskers making contact with the stimulus bar at each jitter position for each angle presented (n=5 mice, 5 sessions, **P<0.01, ***P<0.001, Anova).

### Orientation discrimination requires spatial summation across multiple whiskers

Next we probed how mice discriminate the orientation of the stimulus. First, we confirmed the role of whiskers in task performance. Once mice had reached threshold and stable performance on the task, all whiskers were trimmed to stubs (∼2mm length) so that the whiskers could no longer make contact with the stimulus bar (Figure 2A). Trimming all whiskers reduced performance to chance levels (Figure 2A; p < 0.001, n=6 mice), confirming that the mice solved the task through touch and not through any other incidental sensory cue such as audible motor noises.

We hypothesized that shape perception may depend on integration over the whisker pad. Therefore, we next asked whether contact information from multiple whiskers was required for mice to solve the task. To address this directly, we measured performance as we acutely trimmed the whisker array to different numbers or spatial combinations of whiskers (Figure 2A). We hypothesized that mice would require at least two whiskers, specifically in two different whisker rows (i.e., along an arc), to obtain the relevant information on surface orientation. This was based on the reasoning that whiskers in different rows could discriminate positive from negative surface orientations by computing the relative horizontal position of object contact; in contrast, whiskers in the same row would only access the object at nearly the same elevation, and therefore have far less information on surface orientation.

Indeed, when mice were trimmed to a row of four whiskers their performance in the task was significantly impaired, with discriminability dropping to chance levels (Figure 2B,D,E; p < 0.01, n=7 mice, 14 sessions) and never recovered with subsequent trimming. In contrast, when trimmed to an arc of four whiskers, performance was not significantly affected (Figure 2C-E; p = 0.74, n=5 mice, 10 sessions). In fact, even trimming to just two whiskers in an arc left performance largely intact (Figure 2C-E). Importantly, however, trimming to just one whisker abolished performance (Figure 2C-E). These results demonstrate that mice solved the discrimination task specifically by integrating at least two whiskers in an arc; contact information from a single whisker was not sufficient for task performance (Figure 2E). It seems probable that mice with only one whisker could discriminate surface orientation by adopting alternative whisking strategies or by relying on the differing deflection angles of a single remaining whisker. However, these data indicate that when trained with multiple whiskers in this task, the default strategy the mice use is to summate over at least two whiskers along an arc.

### Mice can use the orientation of the stimulus *per se* to solve the task

Shape perception should be invariant to the vantage point or absolute location of the object. Therefore, we next asked whether some trained mice might actually use the orientation of the stimulus bar *per se* to solve the task. For such mice, performance should be unaffected by presenting the oriented stimulus bar in random horizontal locations within the whisking field. For mice that don’t learn an orientation-based rule, but instead rely on a less general strategy – for example, by identifying a set of whisker contacts in a set of absolute spatial positions – presenting the stimulus bar in additional horizontal locations should impair performance.

To test this hypothesis, we jittered the absolute horizontal spatial location of the oriented bar, trial by trial, and monitored the effect on each mouse’s performance. For these experiments the stimulus bar was moved in from the side (‘Lateral presentation’) rather than from the front to preclude the possibility that mice could solve the task while the stimulus bar was still moving towards its final position (Figure 3A). Although these mice were initially trained with the stimulus moving in from the front, all mice quickly acclimatized to the sideways presentation (1-3 days). Once they regained their original levels of performance, we acutely began presenting the stimulus to one of three final positions along the whisking field that engaged different sets of whiskers. All of the three final jitter positions were randomly interleaved in the test session. Importantly, three out of five of the test mice maintained high levels of performance (d-prime>1.5) across all of the jittered final positions, strongly suggesting that these mice solved the task by sensing the stimulus orientation *per se* (Figure 3B, C). The other two mice only showed a drop for the more distant jitter position, and quickly regained their initial performance levels across all three positions within 1-4 days (Figure 3C). The transient drop in performance for these two mice might be explained either by the difficulty in acclimatizing to altered task conditions (multiple final locations of the stimulus), or by the need to re-learn new spatial and not orientation-based rules for the two additional stimulus positions. The latter explanation would require them to solve the task by remembering three unique spatial coding rules, as opposed to a single orientation-based rule. These data show that mice can solve the task with multiple strategies, but that many mice learn a more generalizable orientation-based rule.

To control for whether mice might adaptively adjust their whisking set point when jittering the position of the stimulus, and therefore still be able to solve the task with a spatial rule (in coordinates invariant to the whisking field), we tracked the motion of the whiskers with high speed imaging. Whisker tracking showed that whisking set-point was largely invariant across the different jitter positions (Figure 3D). Furthermore, high speed video analysis showed that different whiskers primarily contacted the stimulus bar at each of the different jitter positions (Figure 3E). These results argue that mice that showed no acute drop in performance when the stimulus bar was presented to different horizontal locations are using different sets of whiskers to determine the orientation of the stimulus bar and solve the task.

### Barrel cortex neurons encode surface orientation through spatial integration over the sensory array

Next, we sought to address the neural basis for surface orientation discrimination. Since our data show that mice must integrate over multiple whiskers to solve the task, we began by recording neural activity in the barrel cortex whose neurons show substantial integration over the whisker array (Brumberg et al., 1996)(Jacob et al., 2008)(Pluta et al., 2017)(Krupa et al., 2004)(Ramirez et al., 2014). Recent work with controlled whisker deflections has revealed that many primary somatosensory barrel cortex (S1) neurons are optimized encoders of directional whisker movements across pairs of whiskers, offering a conceptual basis for orientation selectivity (Laboy-Juárez et al., 2019).

To address the neural coding of object orientation in the barrel cortex, we trained transgenic mice expressing GCaMP6s in excitatory neurons of the cortex on the task with a full whisker pad (camk2-tta; tetO-GCaMP6s mice) (Wekselblatt et al., 2016). Following training we imaged activity in the barrel cortex with volumetric calcium imaging, collecting neural data from approximately 1,000 neurons per recording in 3 planes of ∼800 x 800μm field of view, encompassing several columns of the barrel cortex. As we were interested in the stimulus-evoked activity we analyzed deconvolved calcium activity beginning at the estimated time of first whisker contact and extending to the end of the behavioral response window, just prior to the reward delivery (Figure 4A, see Methods). This period encompassed the majority of the evoked activity, expanding the window did not substantially change the number or distribution of observed responsive cells (Supplemental Figure 4A-B).

**Figure 4.**
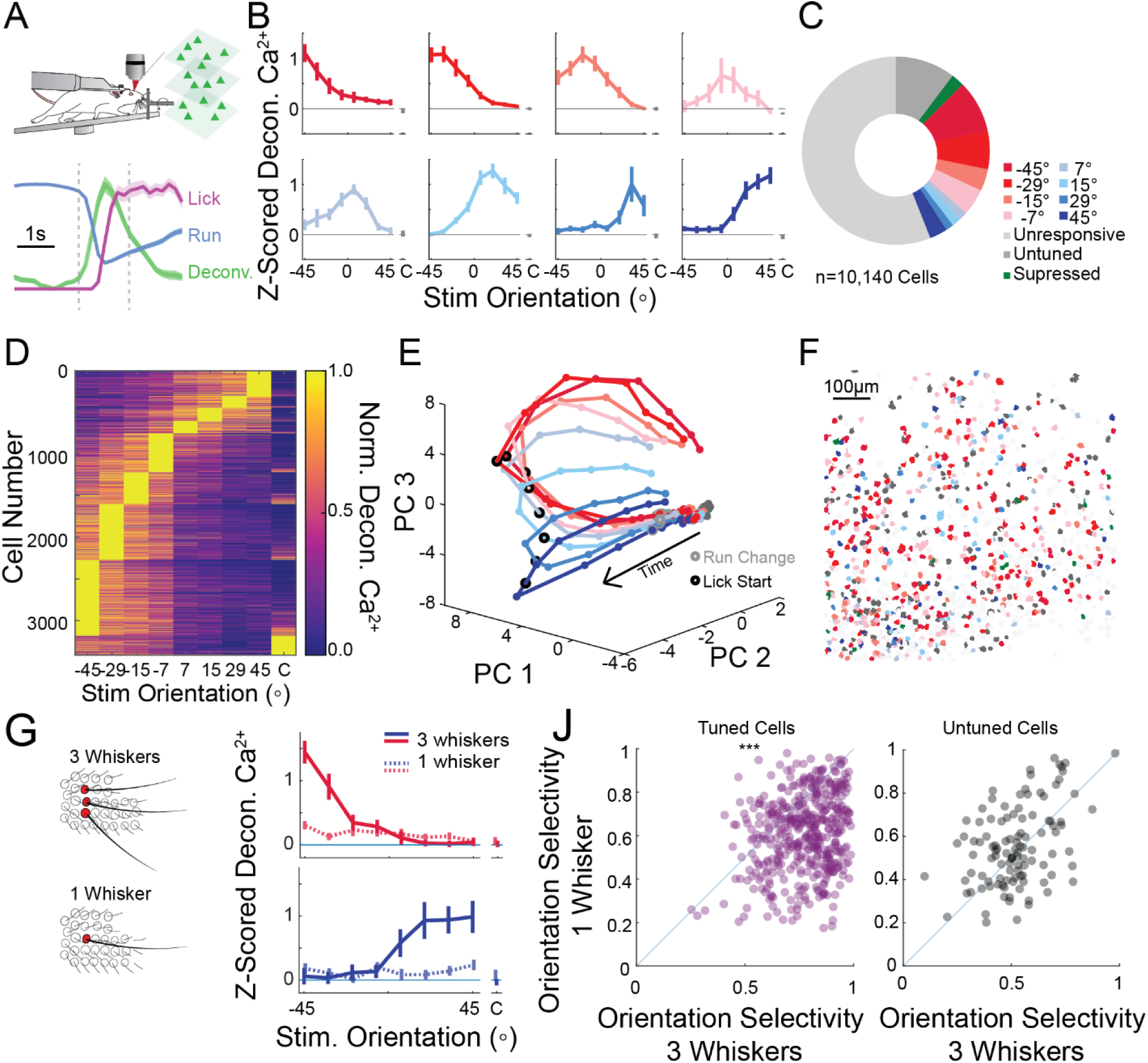
Somatosensory cortical neurons encode each stimulus orientation. A. Schematic of a Ca^2+^ imaging experiment. GCaMP6s-expressing mice are imaged with a two photon microscope while performing the orientation discrimination task. The analysis period is demarcated by the vertical dashed lines (see Methods) and defined by the time window between the initial decrease in run speed and the end of the response window. Normalized mean ± sem for run speed (blue), lick rate (magenta), and deconvolved Ca^2+^ response (all neurons, green) are overlaid to show the relative timing of the response window. B. Mean z-scored deconvolved Ca^2+^ responses of 8 representative neurons from the same mouse, tuned to each of the presented stimuli. Represented as Mean +/-95% Confidence Interval. C. Pie chart showing the relative fractions of all imaged neurons that prefer each stimulus orientation (n=10,140 cells, 9 recordings, 4 mice). D. Tuning curves for all significantly tuned cells (Anova p<0.01) detected across all mice normalized and sorted by preferred stimulus. E. Population activity from a representative recording reduced in dimensionality through PCA. The mean trajectory through PCA space for each oriented stimulus is presented. Each dot is an imaging frame, and the time of run deviation (grey circle) and lick onset (black circle) are noted. F. Stimulus preference map of all neurons recorded in a single recording session; 3 imaging planes are superimposed. Significantly tuned cells are color coded by their preferred orientation (red to blue), untuned but touch-responsive cells are dark grey, and unresponsive cells are light grey. G. Left: schematic of the trimming experiment. Right: orientation tuning curves for two representative neurons during presentation of stimuli with 3 intact whiskers in an arc (solid line), and a single remaining whisker (lighter dotted line). H. Scatter plot of orientation selectivity of neurons before (three whiskers) or after trimming to one whisker. Magenta (*left*), cells that were ‘tuned’ in the three whisker condition and responsive in the one whisker condition (p<3e-41 signed rank test) and grey (*right*), cells that were ‘untuned’ (aka touch responsive but not tuned) before trimming and responsive (with any tuning) in the one whisker condition (p=0.32).

Across the imaged population of neurons, about 45% of identified neurons showed significant responses during presentation of the stimulus bar (see classification in Methods), and about one third showed significant tuning across the stimuli (n=10,140 cells, 9 sessions, 4 mice Figure 4B-D). Importantly, we could identify neurons within each mouse that responded best to each of the eight presented stimuli (Supplemental Figure 4, 5). To quantify the selectivity of neural responses we created a selectivity index (see Methods). Across all cells the selectivity index was much higher than shuffled controls or randomly generated data (p<1e-99, rank sum test, Supplemental Figure 4C). Across the population of imaged neurons we observed a bias towards neurons encoding ‘NOGO’ angles, which may be due to the earlier contact times with NOGO stimuli, more contacts on NOGO trials, or other asymmetries in the stimulation or recording setup (Figure 4C-D and Supplemental Figure 1D). Despite this overrepresentation of cells selective for ‘NOGO’ angles, dimensionality reduction (see Methods) revealed that the population of imaged neurons smoothly represented and could discriminate all stimulus orientations (Figure 4E). In each recording, the population responses for each stimulus orientation diverged shortly after the estimated first whisker contact, indicating a high level of discriminability of the stimulus orientation within the earliest touches (Supplemental Figure 4D). Although the barrel cortex exhibits topographic maps of whisker input (Woolsey and Van der Loos, 1970) and spatial input in the whisking field (Pluta et al., 2017), we did not observe any clear organization of orientation selectivity. Neurons of different orientation preferences were spatially intermingled similar to what is observed in the rodent visual cortex and rodent barrel cortex for passive deflections (Ohki et al., 2005)(Kwon et al., 2018), although distinct for the organization observed for directional tuning to global motion (Vilarchao et al., 2018) (Figure 4F and Supplemental Figure 5).

**Figure 5.**
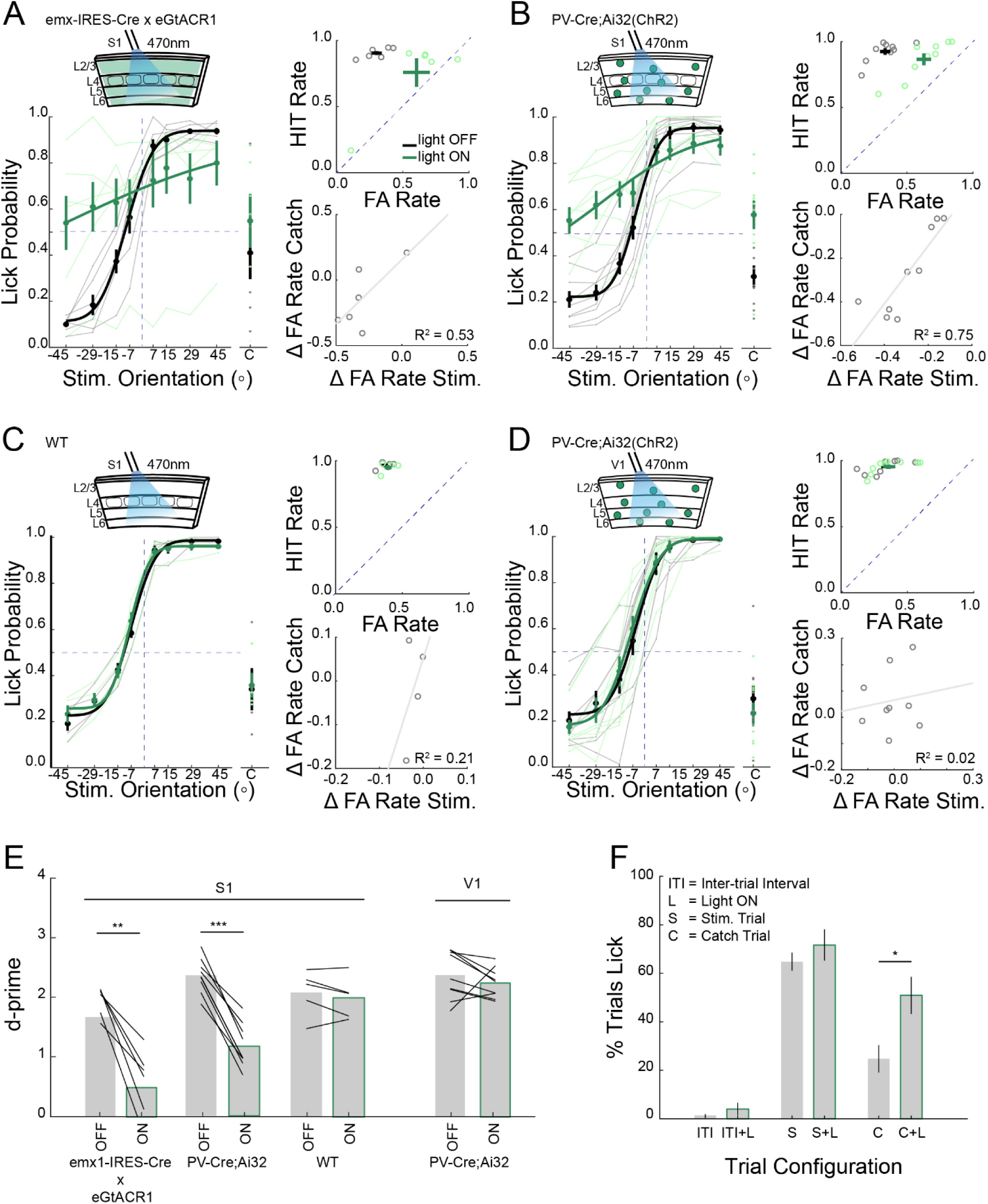
Acute inactivation of the barrel cortex impairs performance in the orientation discrimination task. A. Left top: Schematic of optogenetic suppression of S1. Left bottom: Performance of emx1-IRES-Cre mice virally expressing eGtACR1 on control (black) and optogenetic silencing trials (green). Thick lines: average performance across all mice; thin lines: average performance of each individual mouse. Right Top: ‘FA’ verses ‘Hit’ rate for each mouse, dashed line represents unity line. Right bottom: difference between FA rate for stimulus trials verses catch trials during light ON and light OFF trials (all mice; grey circles, line - linear regression fit, n=6 mice). B. As in A but for PV-Cre;Ai32(ChR2) in S1 (n=9 mice). C. As in A, B but for wild type mice in S1 (n=4 mice). D. As in A-C but for PV-Cre;Ai32(ChR2) mice where the optogenetic illumination was targeted to the primary visual cortex (V1) rather than the barrel cortex (n=9 mice). E. Bar graph of d-prime values for each cohort, black lines: individual mice, bars: mean ± sem. ** p<0.01, ***p<0.001, rank sum test. F. Quantification of licking of PV-Cre;Ai32 mice with optogenetic silencing over S1 during the inter-trial interval (ITI), stimulus trial (S) or catch trial (C) with and without light on (L) (For S and C conditions; n=11 mice, 22 sessions, for ITI conditions 3 mice, 5 sessions, *p<0.05, rank sum test).

To determine whether spatial summation across the whisker array was necessary for the orientation selectivity of the neurons, we acutely trimmed each mouse’s whiskers from an arc of 3 whiskers to a single whisker, a condition in which the mice fail to perform the task (see Figure 2). As a control, orientation tuned neurons were observed at similar rates and distributions in mice with three whiskers in an arc as compared to mice with all whiskers intact (37% tuned, Figure 4G, Supplemental Figure 6A-C, n=2,138 cells, 2 recordings, 2 mice). However, when trimmed from three down to one whisker, we observed roughly half as many tuned cells, but an increase in the proportion of responsive but untuned cells (20% remained tuned after trimming, Supplemental Figure 6A-C). Indeed, among the neurons that were both responsive in the three-whisker condition and retained stimulus-driven activity following trimming, the majority of tuned cells showed a decrease or complete loss of orientation selectivity (p <0.001 1e-41 rank sum test), whereas untuned cells remained untuned (p>0.32 rank sum test, Figure 4J and Supplemental Figure 4C). Further trimming to remove the last remaining whisker reduced task evoked activity to chance levels (Supplemental 6A, B). This implies that integrating across input from two or more whiskers is required for much of the surface orientation coding in the barrel cortex in this task. This across-whisker summation might be occurring in the barrel cortex itself, or may occur upstream, such as in the thalamus, where neurons can also show multi-whisker receptive fields (Timofeeva et al., 2004). Although barrel cortex neurons are well-known to encode angular deflections of single whiskers (Bruno et al., 2003)(Simons and Carvell, 1989), angular tuning does not appear to be sufficient, at least in this task, for much of the orientation coding or behavioral performance. Instead, these data implicate more global computations across the whisker array.

**Figure 6:**
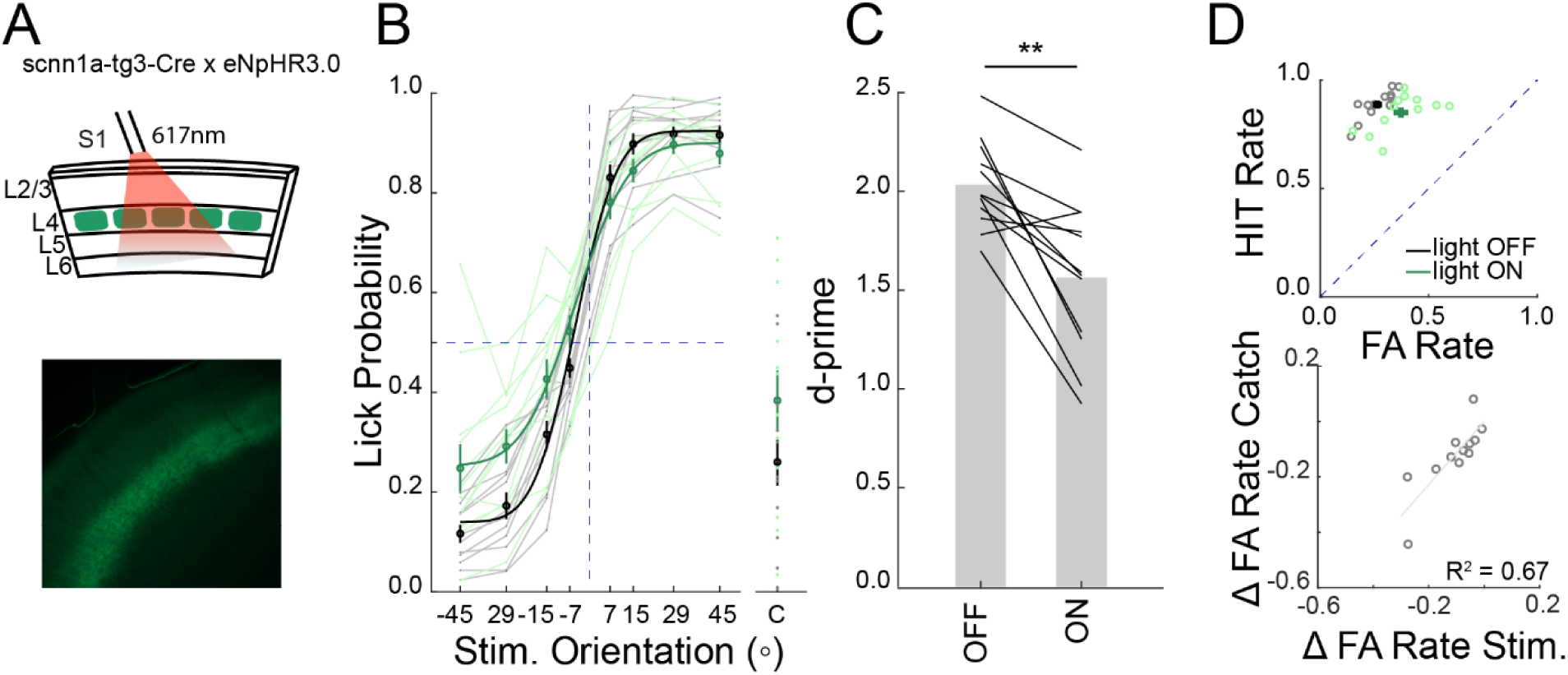
Selective optogenetic suppression of layer 4 excitatory neurons decreases performance in the object discrimination task. A. Top: Schematic of optogenetic suppression of layer 4 excitatory neurons in S1. Bottom: Coronal section through barrel cortex showing eNpHR3.0-YFP expression. B. Performance of scnn1a-tg3-Cre mice virally expressing eNpHR3.0 on control (black) and optogenetic silencing trials (green). Thick lines: average performance across all mice; thin lines: average performance of each individual mice (n=12 mice). C. Bar graph of d-prime values for light off and light on conditions (black lines; individual mice, bars: mean ± sem p<0.01, rank sum test). D. Top: FA rates and HIT rates with and without optogenetic suppression of L4. Bottom: difference between FA rate for stimulus trials verses catch trials during light ON and light OFF trials (all mice; grey circles, line – linear regression fit, n=12 mice).

### Orientation discrimination depends on neural activity in the barrel cortex

Our data above show that mice learn to solve the orientation discrimination task by integrating touch information from multiple whiskers. Although neurons in the whisker system as early as the trigeminal nucleus show sensitivity to multiple whiskers (Minnery and Simons, 2003), prior work (Brecht et al., 2003)(Brumberg et al., 1996)(Pluta et al., 2017) and our imaging data above show that spatial integration is pronounced at the level of S1. Therefore, we tested whether neural activity in S1 would be required for task performance. Since prior studies have shown that the barrel cortex can be dispensable for tactile tasks (Hong et al., 2018)(Hutson and Masterton, 1986), we inactivated S1 neural activity in three different ways. First, we directly suppressed excitatory neurons in the barrel cortex by virally expressing the potent inhibitory opsin eGtACR1 in emx1-IRES-Cre mice (Mardinly et al., 2018). An optic fiber coupled to a 470nm LED was positioned over S1 to illuminate all or nearly all of the barrel cortex. We randomly switched the LED on during one third of all stimulus trials; the LED was turned on for one second preceding, and one second during the response window to ensure optogenetic silencing occurred before the first whisker contact for all stimuli, and throughout the sampling and response window. Optogenetically suppressing excitatory neurons strongly reduced performance on the discrimination task (Figure 5A, E). Next, we photo-stimulated parvalbumin-positive (PV) GABAergic interneurons (PV-Cre;Ai32) to silence nearly all barrel cortex activity by driving widespread inhibition, as many other studies have done (Sachidhanandam et al., 2013)(Guo et al., 2014). Illumination of the barrel cortex in these mice likewise significantly reduced performance (Figure 5B,E). As a control, illumination in mice not expressing any opsin, or targeted inactivation of the primary visual cortex (V1), had no impact on behavioral performance (Figure 5C-E). These results imply that barrel cortex activity is required for surface orientation discrimination. We observed that much of the degradation of performance was due to a strong increase in false alarm rates on ‘NOGO’ stimuli (Figure 5A,B). This could be because mice had a strong bias to lick across all stimuli given how they were trained on the task (Supplemental Figure 1), or because suppression of the barrel cortex directly induced licking behavior. To address this issue, we first analyzed the impact of cortical optogenetic suppression on ‘catch’ trials, when the stimulus was not within reach of the whiskers but the motor carrying the stimulus did move, generating an audible sound. On these trials barrel cortex suppression likewise strongly increased the probability that mice would lick even though there was no contact between the whiskers and the stimulus (Figure 5A,B). Perhaps more importantly, the light-dependent increase in the false alarm rate on catch trials predicted a corresponding increase in false alarms for ‘NOGO’ stimuli during optogenetic suppression but not in control mice (Figure 5A-D). This suggests that eliminating barrel cortex activity may have reduced the ability of mice not only to discriminate stimulus orientation but also reduced their ability to determine whether they were touching a stimulus at all. Next, we silenced the barrel cortex during inter-trial intervals when there was no audible cue that a trial had been initiated (unlike ‘catch’ trials when the stimulus did move but did not reach the whisking field). In contract to ‘catch’ trials, during the inter-trial interval illumination of the barrel cortex did not increase the probability of licking (Figure 5F). This shows that suppressing the barrel cortex did not directly induce the mice to lick in the absence of a contextual cue indicating the initiation of a trial. Rather, these data suggest that the ‘GO’/’NOGO’ training paradigm we employed may have generated a strong bias to lick on trials when mice either could not discriminate stimulus orientation or could not determine whether a stimulus was even present or not.

Although these results imply that barrel cortex activity is required for task performance, they provide no information on which sub-classes of cortical neurons might be important. Next, we specifically suppressed activity in a subset (∼40%) of layer 4 (L4) excitatory neurons with the neuronal silencer eNpHR3.0 in scnn1a-tg3-Cre mice targeted to the barrel cortex (Figure 6A). Suppressing L4 excitatory neurons is a more subtle perturbation than cortex wide inactivation; as we have previously shown, L4 suppression under identical conditions significantly degrades spatial representations in S1, but does not entirely abolish sensory-evoked activity in any cortical layer (Pluta et al., 2015). We found that optogenetic suppression of a subset of L4 neurons significantly impaired performance, albeit more modestly than for total cortical inactivation (Figure 6B-D). This demonstrates that L4 activity is required for normal behavioral performance, mostly likely either by driving sensory input to L2/3, or by sculpting activity in L5 through translaminar inhibition. Taken together, these results imply that barrel cortex activity, *per se*, is necessary for surface orientation discrimination.

## Discussion

Using a novel active tactile discrimination task, high-speed behavioral analysis, optogenetics, and two photon calcium imaging we addressed how animals use active touch to discriminate surface orientation, a process that may be critical for determining an object’s shape and ultimately its identity. Judging shape, whether by vision or somatosensation, requires synthesizing information gathered from different parts of the sensor array into a coherent percept. By taking advantage of the ability to acutely trim whiskers, we determined that mice required summating over multiple whiskers, specifically at least two in an arc, to discriminate the orientation of a stimulus. As barrel cortex neurons are well known to integrate broadly over the whisker array, we found that many barrel cortex neurons were selective for specific orientations, possibly by summating multiple whisker inputs in specific ways. Based on our collected data, we propose a model where active contact of as few as two whiskers with a surface drives sufficient activity in orientation-selective neurons of the barrel cortex to give rise to a percept of a specific surface orientation. More generally, we propose that during natural object exploration, when many whiskers will contact adjacent or overlapping surfaces of an object, spatially distributed orientation-selective activity across many columns of the barrel cortex may give rise downstream to highly shape-specific neurons, akin to those found in primate inferotemporal cortex (Perrett et al., 1982), that might be closely linked to object identification.

Since haptic perception in our task involves active whisker scanning, mice may have solved the discrimination task in a number of ways. One intuitive solution would be for neurons to compute the relative contact time of each whisker in an arc with the oriented stimulus, information that is known to be encoded at more peripheral levels (Jones et al., 2004)(Leiser and Moxon, 2007) as well as in the cortex (Curtis and Kleinfeld, 2009)(Waiblinger et al., 2015). Positive and negative lags of the relative contact time would thus indicate either positive or negative angles, which in our paradigm indicate either ‘GO’ or ‘NOGO’ behavioral choice.

Alternatively, mice could also have used simple XOR logic across pairs of whiskers to discriminate orientations. Although it seems probable that different mice adopt different strategies, a significant fraction of mice seems to perceive the orientation of the bar *per se* since acutely challenging them with the stimulus bar at different horizontal positions resulted in little to no drop in performance (Figure 3). Since mice could not perform the task with just one whisker, it also seems unlikely that mice simply computed the angle of deflection of each whisker, although with further training they may adopt this strategy. Understanding the exact computation that mediates orientation discrimination should be a fruitful subject of additional work.

Importantly, with multiple approaches, we showed that normal activity in S1 is required for animals to discriminate surface orientation in our task. S1 inactivation could impair performance because S1 activity encodes the surface orientation and passes this information downstream to ultimately generate the orientation-selective percept. Alternatively, S1 inactivation might simply disrupt activity in another brain region, such as the superior colliculus, that is more directly responsible for computing surface orientation and passing this on to decision and motor execution circuits. Previously, focal inactivation or ablation of a specific sensory or motor area can lead to spontaneous recovery. This recovery could be explained by homeostatic rebalancing of activity in another brain region whose activity was disrupted by the lesion, or by the animals relearning the task with auxiliary circuits (Hong et al., 2018)(Otchy et al., 2015). Although these are both very difficult to rule out in our experiments, an equally and perhaps more parsimonious explanation is that S1 activity and sensory computation in S1 *per se* are likely to be required. This is supported by the fact that S1 neurons encode the surface orientations explicitly. Finally, we found that degrading spatial representations in S1 via a much more subtle and partial L4-inactivation also impaired performance on the task, demonstrating that normal L4 activity is required, and lending additional support to the notion that S1 computation – putatively via L4 neurons, is necessary for surface orientation discrimination.

Understanding how animals employ active sensorimotor strategies to optimize sensation and generate coherent percepts of the external world is critical for understanding many aspects of brain function and behavior. The head-fixed training paradigm we developed is simple but powerful – learning is quick and robust and can be readily probed by high-speed whisker tracking and neural activity measurement and perturbation. Future experiments aimed at identifying the downstream circuits in the somatosensory system responsible for integrating multiple surface orientations into a specific percept of a 3D object, will yield key insights into high level perceptual processes. Mice offer the most powerful genetic toolkit of any mammalian species, and like other rodents, presumably use high resolution touch input to discriminate and identify small objects near their head. Therefore, mice trained on this and similar tactile tasks might yield fundamental insights into the neural mechanisms of higher order shape perception.

## Acknowledgements

This work was supported by The New York Stem Cell Foundation and by grants from the Arnold and Mabel Beckman Foundation, NINDS grant DP2NS087725-01, the McKnight Foundation, the Simons Foundation Collaboration for the Global Brain award 415569 to I.A.O., and NEI grant K99 EY029758-01 to I.A.O. We thank Dan Feldman and members of the Adesnik lab for a critical reading of the manuscript.

## Author contributions

J.B. and H.A. conceived the study. J.B. conducted all the behavioral experiments. J.B. and I.A.O. conducted the imaging experiments. G.I.T. provided software for whisker tracking. S.G., V.J., and M.V. assisted in behavioral training and experiments. H.A., J.B., and I.A.O. wrote the paper.

## Methods

All experiments were performed in accordance with the guidelines and regulations of the Animal Care and Use Committee of the University of California, Berkeley.

### Transgenic mice

Mice used for experiments in this study were either wild type (ICR white strain, Charles River), PV-IRES-Cre (Jax stock# 008069) crossed with Rosa-LSL-ChR2 (PV-Cre;Ai32) (Jax stock# 012569), emx1-IRES-Cre (JAX stock# 005628), scnn1a-tg3-Cre (Jax stock# 009613), tetO-GCaMP6s (Jax stock # 024742), and/or Camk2a-tTA (Jax stock# 003010). Mice were housed in cohorts of five or fewer in a reverse light:dark cycle of 12:12 hours, with experiments occurring during the dark phase.

### Headpost surgery

For head fixation during behavioral and physiological experiments, a small custom stainless-steel headplate was surgically implanted. Briefly, adult mice (P35-P50) were anesthetized with 2-3% isoflurane and mounted in a stereotaxic apparatus. Body temperature was monitored and maintained at 37ºC. The scalp was removed, the fascia retracted, and the skull lightly scored with a drill bit. Vetbond was applied to the skull surface, and the headplate was fixed to the skull with dental cement (Metabond). A fine-point marker was used to note the approximate location of bregma and the left primary Somatosensory barrel field (S1; 3.5mm Lateral, 1.5mm posterior to bregma) or left primary Visual Cortex (V1; 1mm Lateral, 0mm posterior to lambda), to guide the placement of optical fibers above the skull during optogenetic manipulations. Mice received buprenorphine and meloxicam for pain management and were allowed to recover for at least three days before being placed on water restriction.

### Cranial window surgery

For imaging experiments, a second similar surgery was performed to implant a cranial window over S1. Briefly, mice were anesthetized as above, a 3-3.5mm region of skull located by stereotaxic coordinates (3.5mm Lateral, 1.5mm Posterior to bregma) was removed using a dental drill (Foredom) with a 0.24mm drill bit (George Tiemann & Co.) and/or a biopsy punch (Robbins Instruments). The window was replaced with 3 glass coverslips (two 3mm and one 5mm), and cemented into place with dental cement. Mice were given additional saline during surgery (0.3ml 0.9% NaCl). Mice were not water deprived at the time of surgery, and were given several days to recover before water deprivation resumed. Mice received Buprenorphine and Meloxicam for pain management and Dexamethasone to reduce brain swelling.

### Viral infection

Neonatal emx-IRES-Cre (P3-4) or scnn1a-tg3-Cre mice (P4-5) were injected intracranially with ∼150-200nl of AAV9-CAG-DIO-nls-mRuby3-IRES-ST-eGtACR1 (eGtACR1) or AAV9-EFla-DIO-eNPHR3-EYFP (eNPHR3) respectively at three locations in S1 at approximate locations relative to the lambda suture: AP 1.8-2mm, ML −2.5-3mm. Neonates were briefly cryo-anesthetized and placed in a head mold. Viruses were acquired from or custom produced at the University of Pennsylvania Vector Core.

### Water restriction

Initial animal weight was recorded for 3 days before water restriction to establish baseline weight. Mice were then placed on controlled water and received 1.0 ml/day of water. On training days, mice received the majority of water during the task. Mice were weighed after training and given additional water if their weight had dropped below 75% initial weight. Food was available *ad libitum*. The weight and health (quality of the fur and nails, gait, posture) of the mice were monitored daily.

### Behavioral apparatus

The behavioral apparatus was controlled in real time by an Arduino Mega 2560 or Due, which interfaced with custom written software in Java. Mouse running velocity was measured via an incremental encoder (US Digital). Mouse licking was detected with a custom 2-transistor circuit between the 0.05-inch diameter steel tube lickport and a stainless steel headpost of the mouse (Slotnick, 2009). Whisker contacts were detected using a custom designed sensor attached to a 3D printed stimulus bar emitting an infrared (IR) beam that when broken by a whisker was recorded by the IR photodiode as a voltage deflection. Water was delivered by gravity through a lickport under solenoid valve control (Neptube Research Inc.). A linear bar (25 x 3 x 1 mm) was coupled to a stepper motor (NEMA 8: “stimulus motor”) via a custom cylindrical arm. When in the ‘home position’, the stimulus motor rotated the linear bar clockwise in the vertical (dorsal-ventral) direction between 360° to 720° to select a new orientation each trial. The stimulus motor was mounted at a 90° to a larger “position motor” (NEMA 17). Once the mouse reached the required run threshold to initiate a new trial, the position motor rotated counter-clockwise until it reached the target area (pre-defined by an IR-beam break), whereby the motor paused for 1 second during the response window, before moving backwards out of the whisker field until it reached the home position. The trajectory of the stimulus was aligned in roughly the same horizontal plane as the C row whiskers of the mouse’s right whisker field, and centered around whisker C2. The distance between the stimulus and the whisker pad was 10.5 ± 0.3mm from follicle pad, n=10. For jitter experiments, stimulus bar moved in laterally from the mouse whisker pad. Stimulus bar was jittered 20.6 ± 1.2 degrees for first contact and 39.1 ± 0.9 degrees at its final resting position relative to the mouse’s whisker pad. The entire apparatus was mounted on an 8.0 x 8.0 x 0.5 inch (Thorlabs) anodized aluminum breadboard and enclosed in a light isolation box (80/20). Negative reinforcement was delivered via a single puff of compressed air, gated by a solenoid valve, to the contralateral eye from the whisker stimulus from a distance of ∼2 mm through a 0.05-inch-diameter steel tube. Air puff pressure was increased until it produced a blink response by the mouse. An optical fiber coupled to an LED was placed on a separate manipulator above the mouse’s head. On sessions with photo-stimulation, the fiber was positioned over the approximate location of the barrel cortex, as close to the thinned skull as possible. A masking LED light to minimize behavioral detection of the optogenetic light was also used on every trial during photo-stimulation sessions. The masking light was positioned directly in front of the mouse’s eyes and would turn on each trial for 1 second preceding and 1 second during the response window.

### Orientation discrimination task

The orientation discrimination task followed the classical ‘GO/NOGO’ paradigm. Head-fixed mice were trained to lick when they detected a ‘GO’ stimulus and withhold licking when they detected a ‘NOGO’ stimulus. Mice were presented with stimuli in one of eight possible orientations, defined by their angular position in the vertical plane. The possible stimulus orientations were ±45°, ±29°, ±15°, or ±7° from dorsal-ventral axis. ‘GO’ stimuli were angled in the posterior direction and arbitrarily defined as positive angles. ‘NOGO’ stimuli were angled in the anterior direction and defined as negative angles. On each trial, stimulus orientations were chosen at random subject to the constraint that all eight stimulus orientations be presented once in each block of eight trials, and no sequence of more than 3 ‘GOs’ or 3 ‘NOGOs’ in a row. On a subset of trials (10%), ‘catch’ trials were randomly presented whereby the stimulus stopped just anterior to mouse and outside of the whisker field.

The sequence for each trial was as follows (Supplemental figure 1). Each trial began with the stimulus in the home position for a 3-5 second waiting period. During this period, the stimulus motor rotated the stimulus to one of the possible pseudo-randomly selected orientations. After the waiting period, mice initiated the start of each trial by exceeding a ∼50 cm running requirement. This triggered the positioning motor to move the stimulus to the target position, within reach of the whiskers, except for ‘catch’ trials where by it was held just outside of the whisker field. Because the target position was in the approximate midpoint of the whisker field, mice typically made their first whisker contact with the stimulus before it reached the target position (Figure 1C, D). Once stationary in the target position, mice were trained to either lick or withhold licking during a 1 second response window. The reinforcement schedule was as follows (Supplemental figure 1: correctly licking for a ‘GO’ angle (‘HITs’) were rewarded with a drop of water (∼5 μl) 0.5 seconds after the response window ended. Incorrectly licking for a ‘NOGO’ angle during the response window (‘False Alarm (FA)) immediately triggered an air puff and a 5 seconds “time-out” period was added to the waiting period at the end of the trial. Correctly withholding licking to ‘NOGO’ responses (“correct rejections”) were not rewarded and incorrectly not licking for ‘GO’ responses (“MISS”) was not punished. All licks were recorded but licks outside of the response window had no consequences. At the end of the response window, the positioning motor moved the stimulus in reverse out of the whisker field and back to the home position. The exact location of the home position randomly jittered each trial by ± 2cm so that the mouse could not predict ‘catch’ trials based on the time in which the motor is moving. Once in the home position a random time out was given if the last trial was a false alarm and then the stimulus motor rotated the stimulus to a new orientation during the waiting period.

### Behavioral training

Training began after headplate application, recovery (∼3days), and 5-10 days of water restriction. We did not find it necessary to handle the mice extensively before training or anesthetize them before head-fixation. Prior to training with the behavioral apparatus, mice were habituated to head-fixation on a free-spinning circular treadmill (diameter, 6 inches) for 4 daily sessions lasting ∼60 minutes each. The training schedule was as follows (Supplemental figure 1): in the first stage of behavioral training (‘1Ori_auto’_), mice were classically conditioned to lick in response to the stimulus. The stimulus was only presented in the ‘GO’ +45° orientation. On every trial, stimulus presentation was paired with a drop of water delivered 0.5 s into the response window. Once mice were reliably licking before the water was delivered (i.e. showing anticipatory licking), they were moved onto the operant conditioning phase of training.

In the second stage of training (‘1Ori’), mice were operantly conditioned to lick in response to the stimulus and withhold licking during the no stimulus, ‘catch’ trials (10% of the trial count). The stimulus was only presented in the ‘GO’ +45° orientation. A water reward was delivered only if mice licked during the response window to the ‘GO’ stimulus. Once mice reliably licked upon detection of the stimulus and withheld licking to catch (>70% correct), they moved to the simplest stage of the discrimination task (‘2Ori’).

During ‘2Ori’ discrimination training, mice were conditioned to lick only for ‘GO’ trials (+45°) and withhold licking to ‘NOGO’ (−45°) stimulus. ‘GO’ and ‘NOGO’ trials were randomly interleaved. ‘Catch’ trials were presented for 10% of trials. A water reward was delivered on ‘GO’ trials (+45° orientation) only if mice licked during the response window. In contrast, an air puff and time out were delivered if mice licked to the ‘NOGO’ stimulus. When mice performed >70% for two consecutive days they were advanced to the final stage of the discrimination task (8Ori). During ‘8Ori’, 4 ‘GO’ and 4 ‘NOGO’ stimuli were randomly presented to the mouse: ±45°, ±29°, ±15° and ±7°, and ‘catch’. Mice had to achieve above 1.5 d-prime during ‘8Ori’ value to be used in subsequent experiments.

Mice were trained 5-7 days per week. All mice had a full pad of whiskers during training. All sessions on ‘2Ori’ and ‘8Ori’ were preceded with ∼10 trials on ‘1Ori_auto_’ to ensure the lickport was positioned correctly and to prime mice for training.

### Whisker tracking

A high-speed camera (Basler acA2000-340km) was placed below the running wheel; whiskers were imaged from below using a telecentric lens (Edmund Optics NT58-257) and a mirror angled at approximately 45 degrees. Some mice and experimental setups required slight adjustments to the mirror angle to properly view the whiskers and object interaction. Whiskers were backlit from above using high-powered diffused infrared LEDs (CMVision-IR200). High-speed videos were acquired with a framegrabber (Silicon Software) at 500 frames per second with a 100-μs exposure and were synchronized with behavioral data via external triggers. Whisker tracking was performed offline using Whisk (Janelia Farms, Howard Hughes Medical Institute), which returned whisker angles and positions for every frame. Tracking data was further processed and analyzed using custom MATLAB scripts written to extract set-point, amplitude, phase, and frequency. Briefly, set-point was calculated as the average of the upper and lower peak envelope of the angle signal, amplitude was the difference of the upper and lower peak envelope, phase was the Hilbert transform of the signal, and frequency was the number of complete whisk cycles for a given unit of time. For naïve vs. expert mouse analysis (Supplemental figure 3), whisker set point was analyzed during the 300ms period following a 2 standard deviation change in baseline run velocity as a proxy for first whisker contact. For analysis of whisker set point during jitter experiments (Figure 3), the period between the change in run velocity (2 standard deviations from baseline) to the onset of lick response was used. Baseline whisking was calculated between –1.0 and –0.7 second before the whiskers had contacted the stimulus bar. A subset of videos were manually annotated for either first contact time (Supplemental Figure 2) or to identify which whisker arc primarily contacted the bar. Human obverses were blind to the experimental condition being analyzed.

### Whisker trimming

A subset of mice had their whiskers either abruptly trimmed from full pad to no whiskers, or progressively trimmed from a full pad to three or four adjacent whiskers either in the same row (C1-C4) or arc (B2-D2), then to two adjacent whiskers either in the same row (C1-C2) or arc (B2-C2), then to one whisker (C2), and then no whiskers. Once mice reached ‘8Ori’ criterion, mice were tested for two sessions prior to trimming in order to determine baseline performance. After each subsequent trimming procedure, mice were tested with the resulting configuration of whiskers over two sessions. For the progressive trimming group, mice were randomly assigned to the row or arc whisker conditions. Only whiskers ipsilateral to the stimulus were trimmed in precise configurations. When trimming from a full pad to four whiskers, mice were briefly anesthetized with 2-3% isoflurane. All other trimmings were done while mice were head-fixed and running on a treadmill. For each of the different progressive trimming conditions, mice were tested over two behavioral sessions usually in consecutive days.

### Calcium imaging

Calcium imaging experiments were performed in mice expressing GCaMP6s in excitatory neurons via tetO-GCaMP6s x Camk2a-tTA or intracranial injection of AAV9 syn-GCaMP6s (titer 8e11 vg/mL). After reaching training proficiency, mice were implanted with a cranial window on the side contralateral to stimulus presentation, following recovery they were returned to training for several days before being moved to an identical behavioral training apparatus connected to a 2-photon microscope (Sutter MOM, Sutter inc.). Mice were given an additional 1-3 training days on the microscope rig to acclimate to the new environment. As two photon microscopes provide many additional distractors to trained mice (e.g. temperature, sounds, environment) we reduced our d-prime threshold for inclusion to 1.0 (60% of recordings passed this threshold). Imaging fields of view were identified by manually deflecting whiskers and navigating towards areas with substantial broad GCaMP fluorescence. Subsequent recording days were placed such that the same cells would not be recorded in separate days. All recordings were performed in L2/3 imaging three 800 × 800 µm planes, spaced 30-50µm apart, at 5.2-6.2Hz with ∼100mW 920nm laser light (Coherent Chameleon) using a resonant galvo system. Images were acquired using ScanImage (Vidrio Inc.) with custom behavioral control software. Tiff files were motion corrected, source fluorescence and OASIS deconvolved signal data was extracted using suite2p (Pachitariu et al., 2016).

As GCaMP6s has slow kinetics, we focused our analysis on a period after the animal’s whiskers had made contact with the object but before the animal had made a behavioral response. For each recording, we identified the imaging frame in which the animal began slowing down, which was highly correlated with the first whisking touches (see Supplemental Figure 1), this frame until the end of the ‘behavioral response window’ is called the standard window and was used for subsequent analysis (Supplemental Figure 4A). Every cell’s OASIS deconvolved activity was z-scored across the entire recording, and each trials’ evoked response was determined as the mean response in the analysis period baselined to a pre stimulus period. Only trials in which the animal responded correctly were used. A trial was excluded if its run speed before stimulus presentation deviated more than 2 S.D. from its mean run speed. Tuned cells were defined as cells whose evoked responses to each oriented stimulus passed an Anova p<0.01, these cells were further divided by their peak response. If a cell responded more on the catch trials than any stimulus it was categorized as a suppressed cell. Touch cells were defined as cells that were not tuned but passed an Anova p<0.01 to all orientations and the catch trial. In all Calcium Imaging based figures error bars denote a 95% confidence interval. Principle component analysis (PCA) was performed on the full unrolled deconvolved calcium activity traces of every cell per experiment and subsequent responses were averaged by stimulus condition. Maps of tuning preference are presented as the projection of all three imaging planes with the outline of each detected source color coded by preferred tuning or touch responsive category. Orientation Selectivity is calculated as the Euclidean Norm of the mean response to each oriented stimulus. As it is difficult to be certain what 0 activity looks like in calcium data this response vector (v) was normalized from minimum to maximum activity. A selectivity of 1 would be a response to only one orientation, whereas a selectivity of 0 would be the same response to all stimuli.

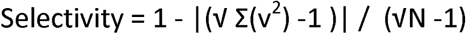

In trimming experiments, intrinsic imaging was first performed to identify which whisker was optimally located in the cranial window for imaging. Whiskers were then trimmed to a column and the experiment began as described above. Roughly half-way through a typical recording session, the whiskers above and below the principle imaged whisker were removed to create the single whisker recording set. As mice are unable to perform the task with a single whisker, recordings are included only if the animal had an performance accuracy d-prime of 1.0 or higher on the 3 whisker condition, but all trials (correct and failures) are included in analyses.

### Intrinsic imaging

Intrinsic optical imaging was performed through the cranial window to localize the C1 and C2 barrels. Prior to imaging, anesthesia was induced as described above and then the mice were administered 0.01 mg/kg Xylazine. Anesthesia was maintained with 1% isoflurane during imaging. Imaging and stimulation were conducted using custom software written in MATLAB. Mice were given 24 hours to recover.

### Optogenetic stimulation *in vivo*

For optogenetic silencing of neurons via eGtACR1 in emx1-IRES-Cre mice or activation of PV interneurons in PV-Cre;Ai32 mice, we used blue light (470nm, fiber coupled LED, Thorlabs) at 10-11 mW/mm2 and to activate eNpHR3.0 in scnn1a-tg3-Cre mice we used 617nm LED at 15-18mW/mm from the end of a 400µm diameter optical fiber. Light intensity was controlled by analog outputs to the LED driver and calibrated with a photodiode and power meter (Thorlabs). The fiber was placed as close to the thinned skull over the S1 region as possible. For behavioral experiments, a square light pulse was applied for 2 second intervals. To ensure photoinhibition before the first whisker contact and throughout the response window, the photo-stimulus started 1 second prior to the stimulus reaching the target position and was sustained until the end of the response window. Photo-stimulation trials were a randomly chosen 33% of all trials. A masking light (blue for eGtACR1 and PV-Cre;Ai32 experiments, red for eNpHR3.0 experiments) was used to control for LED stimulation on all trials and an eye patch was positioned over the right eye of the mouse to prevent any visual cue which may have been gained through the masking light.

### Histology and image acquisition

To quantify the level of expression of the transfected opsins in cortical L5 neurons, mice were perfused and their brains were sectioned and examined under a fluorescent microscope. Immediately after electrophysiology experiments, mice were anesthetized with 5% isoflurane and ketamine and perfused transcardially with 4% paraformaldehyde (PFA). Their brains were removed, stored overnight in 4% PFA at 4°C, and then cryoprotected for at least 24 hours in 30% sucrose buffer solution at 4°C. A sliding microtome (American Optical Company) was used to take 40 μm coronal sections of the barrel cortex and POm-region of the thalamus. Sections were mounted on glass microscope slides using Vectashield mounting medium with DAPI to non-selectively stain cells and protect tissue from photobleaching. Regions of some sections were imaged using a fluorescent microscope (Olympus) coupled to a digital camera (QImaging). To simultaneously view fluorescence from eArch3.0 or eNpHR3.0 expression, DiI, and DAPI, images of a single section were taken with separate excitation filters and merged in Matlab (Mathworks).

### Behavioral data analysis

Although percent correct was used during training to determine criterion, this is more sensitive to response bias, such as tendency to disengage and stop responding for blocks of trials, compared to other measures (Macmillan and Creelman, 1991). For this reason, the discriminability index (‘d-prime’) was used to more precisely evaluate the ability of the mice to discriminate between ‘GO’ and ‘NOGO’ stimulus orientations. In calculating d-prime, only the hit rate (HIT Rate = Hit/Hit+Miss) and false alarm rate (FA Rate = FA/FA+CR) are taken into account. More precisely, d-prime is a measure of the difference between the z-transforms of HIT rate and FA Rate: d = norminv(Hit Rate)-norminv(FA Rate).

Therefore, d-prime is maximized when the subject maximizes Hits (and thus minimizes Miss) and minimizes FA (and thus maximizes the CR), indicating greater ability to discriminate ‘GO’ from ‘NOGO’ stimulus orientations. The effective limit of d-prime is 6.93, typical values are around 2.0, and we have selected 1.5 as a threshold for reliable discrimination. Chance levels of discriminability correspond to d-prime=0. We report overall d-prime values for a complete session as well as d-prime for pairs of ‘GO/NOGO’ stimuli whose orientations were equal and opposite. For the analysis of lick probability, we quantified the proportion of trials where mice made a ‘GO’ response. Psychometric curve of lick probability was fit with Wichmann and Hill psychometric fit.

If a mouse failed to make ‘GO’ responses for 32 consecutive trials, we assumed the mouse was sated and excluded these trials and any subsequent trials from our analysis, except where the mouse had no whiskers or the stimulus was out of reach of the whisker field. If a mouse did not reach 240 trials within a session, data from that entire session was excluded. All times reported during a trial were measured from the onset of the response window (0 seconds).

### Statistics

All analyses were performed using Matlab (Mathworks). The analyses performed were ANOVAs, with multiple comparisons, Wilcoxon rank sum test and Bonferroni test. Unless otherwise noted, all tests were two-tailed and all plots with error bars are reported as mean ± sem. Sample size was not predetermined using power analysis.

---

**Supplemental Figure 1.**
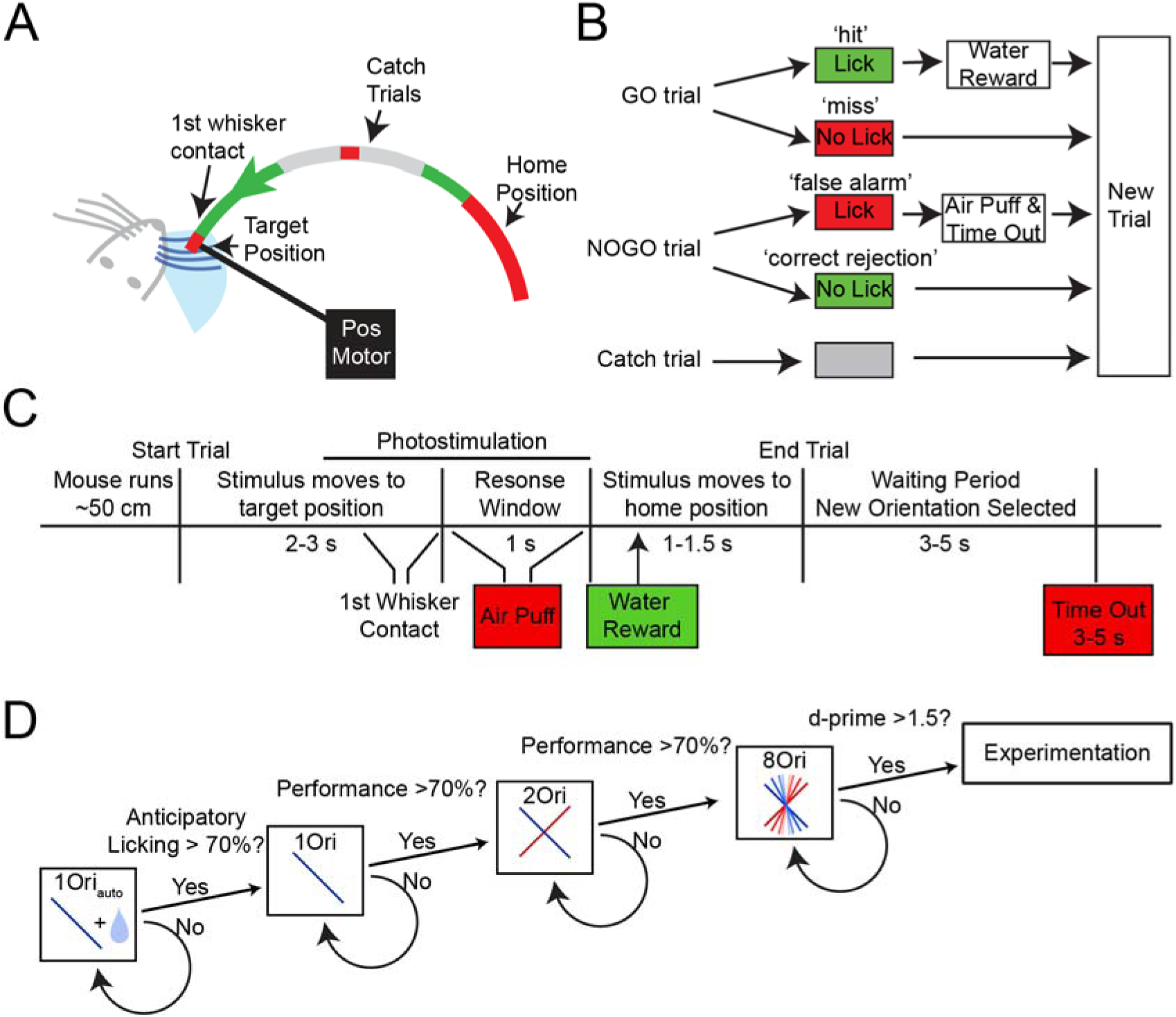
Orientation Discrimination Task Design. A. Schematic of the task. The stimulus bar approaches the mouse from the rostral position on a stepper motor. Mice can make first contact with the bar as it travels into its final resting position within the mouse’s whisker field. For ‘catch’ trials, the stimulus bar is held outside the whisker field. The ‘home position’ is jittered on each trial (red area) so that mice cannot predict ‘catch’ trials based on duration of motor travel. B. Trial structure. For ‘GO’ trials mice receive a water reward for correctly licking during response window (‘hit’). Mice receive an air puff to the eye and a time out if they lick during ‘NOGO’ trials (‘false alarm’). Incorrectly withholding licking for a ‘GO’ trial (‘miss’), or correctly withholding licking for a ‘NOGO’ trial (‘correct rejection’) and catch trials are not punished or rewarded. C. Timeline of task structure. D. Progressive training paradigm. Mice initially are acclimated to the behavior box and running wheel for 4 days. Mice then enter 1 orientation auto ‘1Ori_auto’_ phase of training whereby the ‘GO’ stimulus is presented to the mouse whisker field and automatically paired with a water reward. Once mice show anticipatory licking, the automatic reward is removed and mice only receive a reward if they correctly lick for a ‘GO’ angle ‘1Ori’, 10% catch trials are introduced here too. Once performance reaches >70% mice progress onto the discrimination task by presentation with the −45 ‘NOGO’ angle, then must learn to withhold licking to the ‘NOGO’ and lick for ‘GO’ ‘2Ori’. Once performance exceeds 70%, mice are presented with the full version of the task ‘8Ori’ and must reach d-prime> 1.5 to be used in subsequent experiments.

**Supplemental Figure 2.**
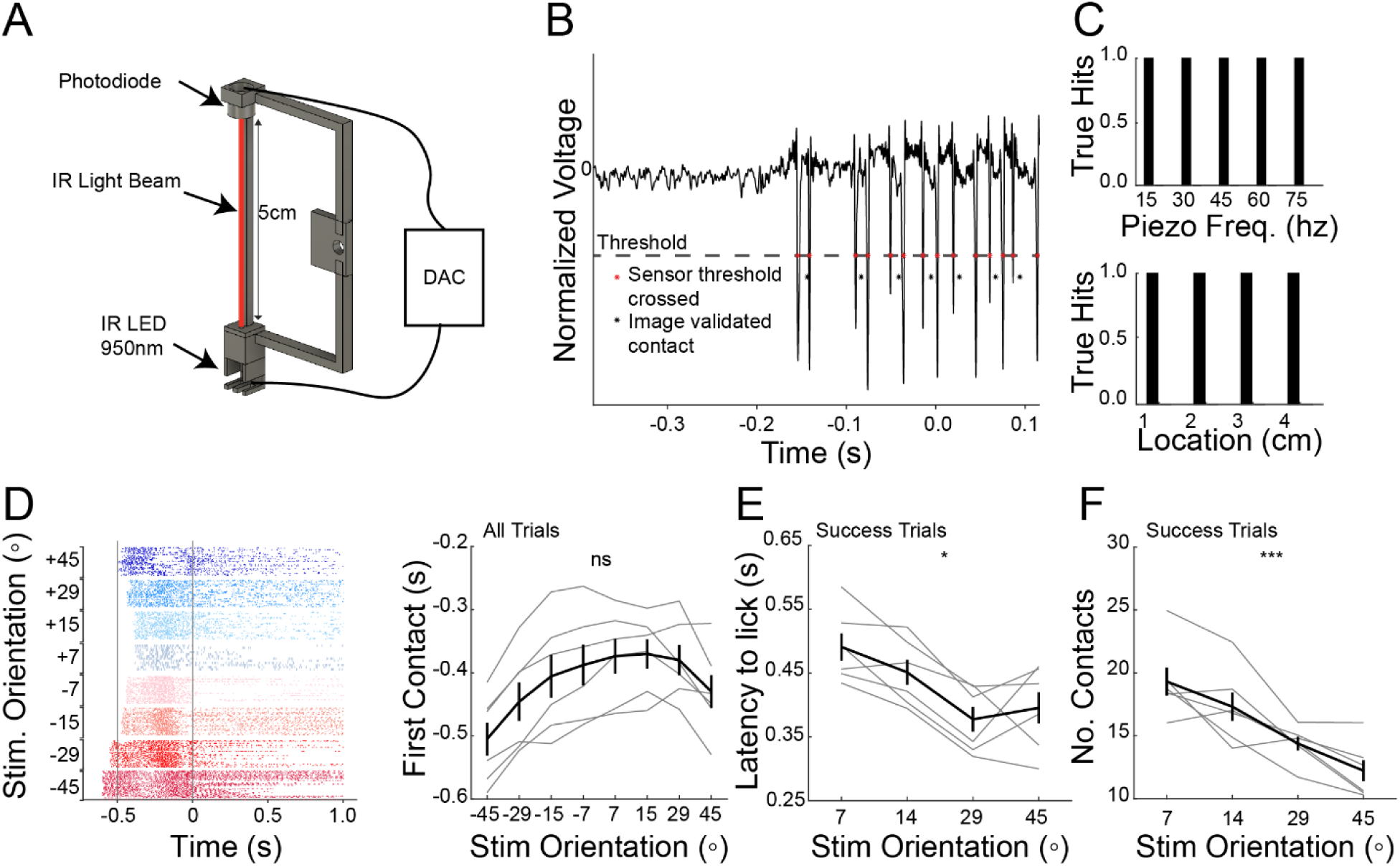
Identification of whisker contacts during the task. A. Schematic of 3D printed stimulus bar with IR LED beam and photodiode for touch detection. B. Validation of the touch detector with simultaneous high-speed imaging. Raw voltage signal from the photodiode; voltage deflections occur when a whisker crosses the IR beam. Grey dotted line indicates the threshold used to identify beam breaks. Red dots indicate detected threshold crossings from the photodiode and black dots indicate visually identified whisker contacts via simultaneous high-speed imaging. Comparison of visually identified contacts from high speed video to putative contacts based on threshold crossings on the photo-diode; accuracy of 94.4% true hits and 11% false positives (n=3 mice, 3 sessions, 30 trials). C. A piezo with a mouse whisker attached was used to further validate the touch detector at different frequencies (top), and at different contact points along the stimulus bar (bottom) (90 trials at each condition). True hits were assessed with simultaneous high-speed imaging. D. Left: Raster plot for an example mouse showing whisker contacts (colored tick marks) sorted by stimulus orientation presented. Grey bars indicate time points –0.5 and 0ms. (n=3 mice, 6 sessions, p>0.05, Anova). E. Latency from first whisker contact to lick response for success trials (n=3 mice, 6 sessions, p<0.01, Anova). F. Number of contacts before lick response on success trials (n=3 mice, 6 sessions, p<0.001, Anova).

**Supplemental Figure 3.**
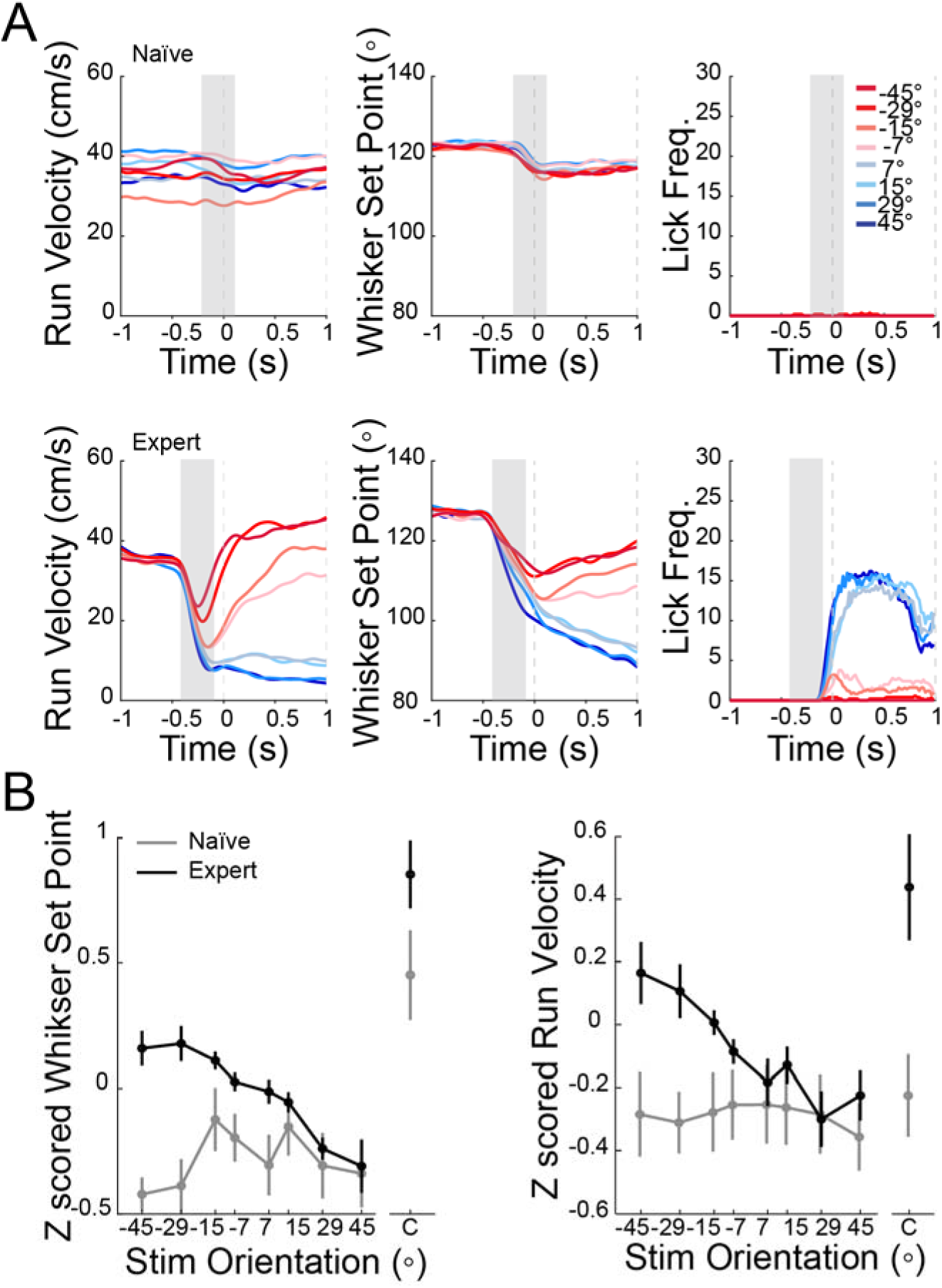
Learning induced changes in running, whisking and licking behavior. A. Run velocity (left), whisker set point (middle) and lick rate (right) for an untrained mouse (top: naïve) and a trained mouse (bottom: expert) presented with the stimulus bar at 8 orientations ‘8Ori’. Colors represent different orientations. Grey shaded area represents running analysis window, dotted lines represent response window. B. Z-scored change in whisker set point (left) and run velocity (right) for naïve (grey; n= 5 mice, 8 session) and expert mice (green; n = 5 mice, 10 sessions) across all orientations (p<0.001, Anova).

**Supplemental Figure 4.**
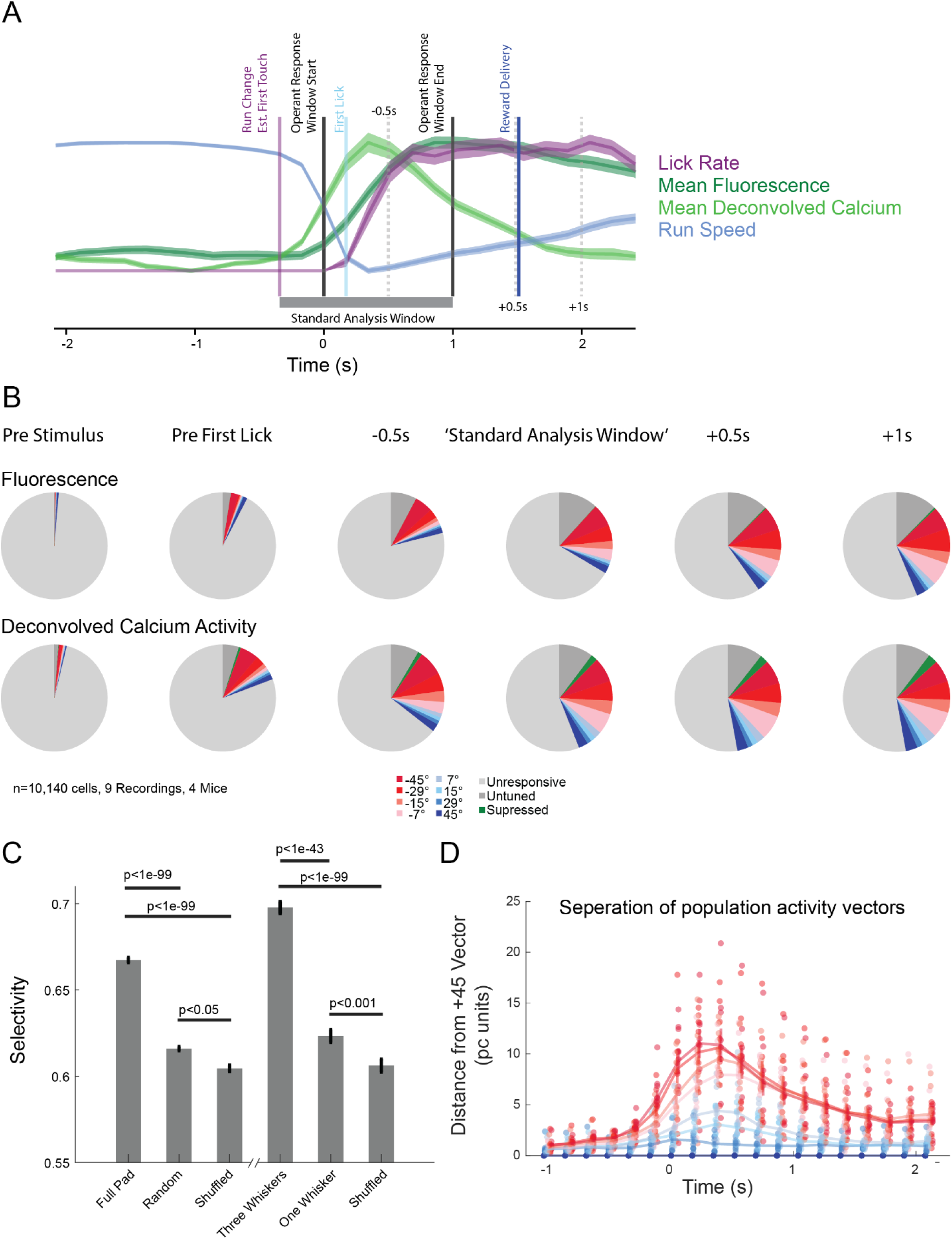
Establishing Window for Calcium Response Analysis. A. Representative trace of a mouse’s run speed (all trial types, blue), Lick Rate (Hit trials, purple), mean calcium fluorescence (mean z-scored dF/F, dark green), or deconvolved calcium response (mean z-scored deconvolved calcium activity, light green). The standard analysis window (grey bar) is defined as the beginning of the deviation in running to the end of the response window. Also marked: The running deviation frame, aka the estimated time of first contact (Purple). The operant response window, when the stimulus is stopped and licks are counted as responses (Black lines). The beginning of the lick response (in hit trials, Light blue). The time of reward delivery (Dark blue). Additionally, three other periods –0.5s, +0.5s or +1.0s from the end of the response window (grey dotted lines). B. The proportion of cells that appear touch responsive or tuned to an orientation is compared as a function of the size of the analysis window (from the estimated first touch to the indicated region), when analyzing z-scored dF/F fluorescence (above) or z-scored deconvolved calcium responses (below). The color (*blue* to *red*) indicates preferred orientation of neurons, *dark grey* indicates untuned neurons, *green* suppressed neurons, and *light grey* unresponsive neurons as in Figure 4. C. Selectivity index computed for all detected neurons from normal experiments (mean ± sem.). compared to similar metrics created by random values or shuffled data. P values are rank sum comparisons between populations. *Right*, selectivity index from neurons from the trimming experiments D. Euclidean distance of population activity across the first 3 principle components. Measured in each frame as the distance from the +45° vector to each of the other angles. Faint dots are each recording’s average, solid line mean ± sem.

**Supplemental Figure 5.**
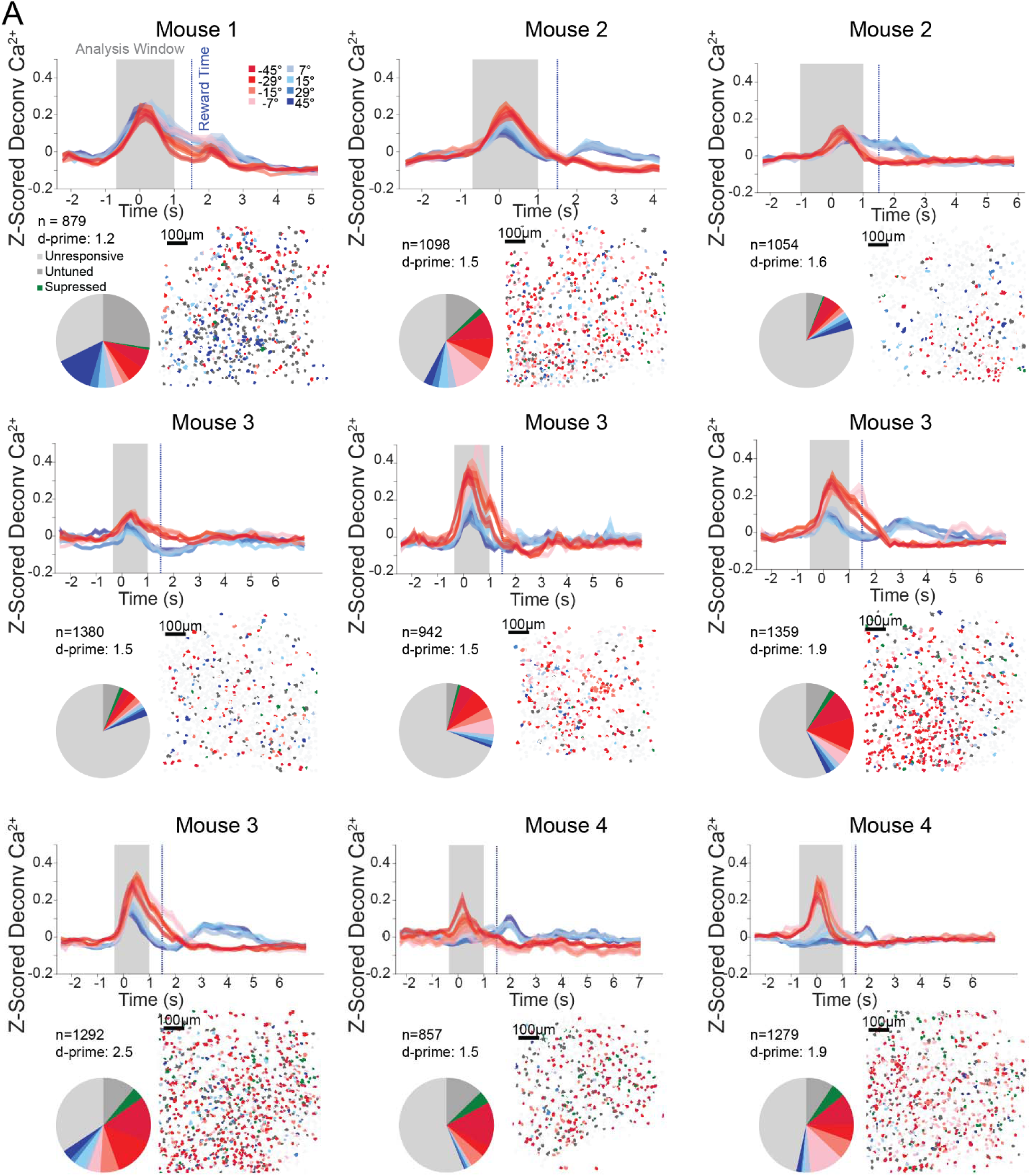
Calcium Responses from Each Field of View. A. For each recording, *top*, mean (±95% confidence interval) population deconvolved calcium activity for each presented angle. *Grey section*, the analysis window used determined by the start of a change in running (a.k.a. estimate first time of licking) until the end of the behavioral response window. *Blue Dotted Line* the time of reward delivery. *Inset:* The Number of detected neurons, and d-prime recorded during this session. *Bottom Left*. Pie chart showing the relative percent of neurons that were unresponsive, untuned, suppressed, or tuned as in Figure 4C. *Bottom Right*, stimulus preference map of all neurons recorded in a single recording session; 3 imaging planes are superimposed as in Figure 4F.

**Supplemental Figure 6.**
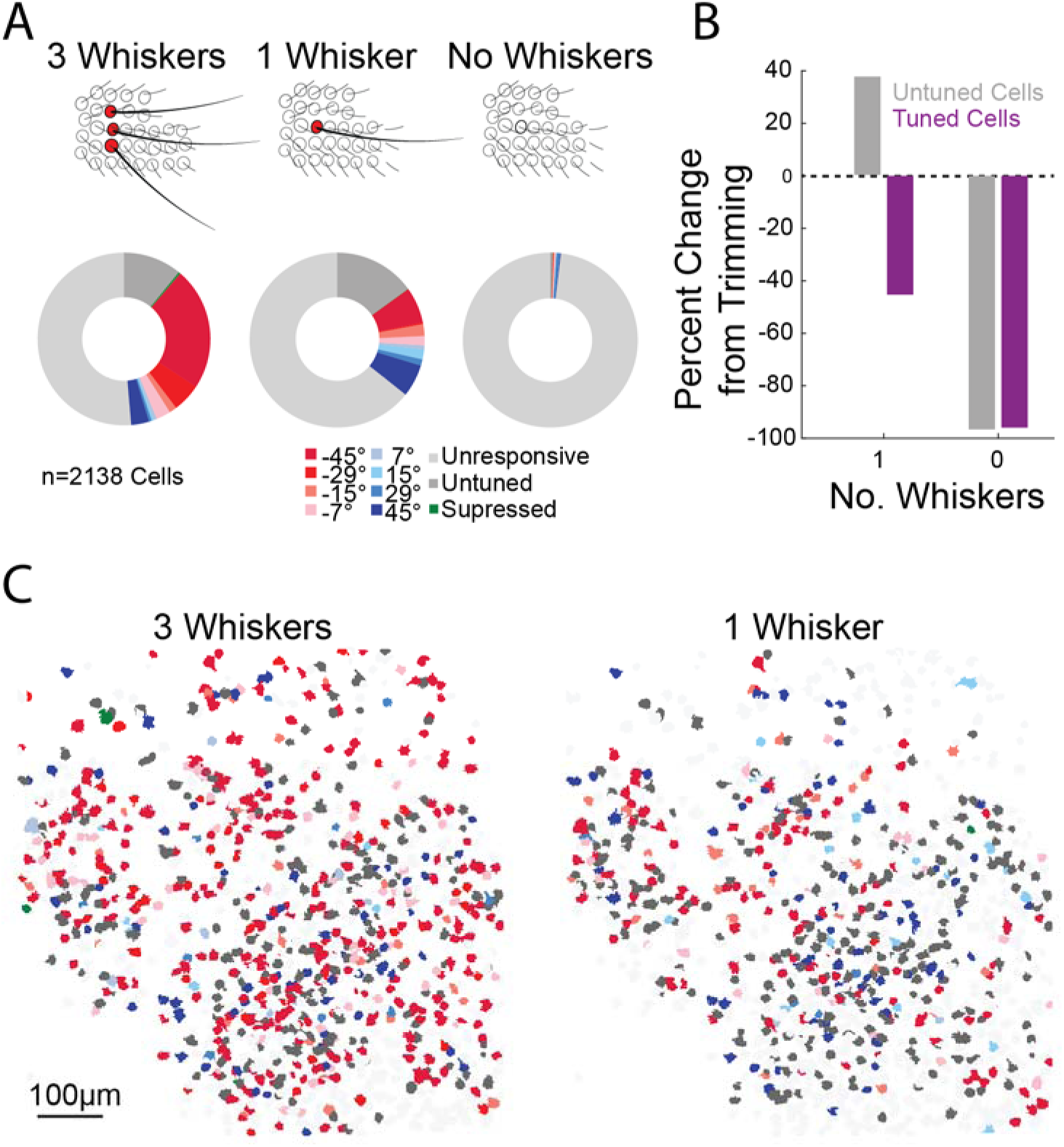
Whisker Trimming reduces the number of Tuned Cells. A. Top: schematic of the trimming experiment. Bottom: Fraction of neurons that are significantly tuned or untuned but touch responsive in the three whisker, one whisker, or no whisker condition. Color Code as in Figure 4C. B. Relative change in the number of cells that could be classified as touch responsive but untuned (‘untuned’, grey) or significantly tuned to any orientation (‘tuned cells’, magenta) for the one and no whisker conditions relative to the three whisker condition. C. Maps of the locations of the recorded neurons (three planes superimposed) colored according to their preferred stimuli, with 3 (left) or 1 (right) whiskers remaining.

